# Cytotoxicity of 1-deoxysphingolipid Unraveled by Genome-wide Genetic Screens and Lipidomics

**DOI:** 10.1101/658690

**Authors:** A. Galih Haribowo, J. Thomas Hannich, Agnès H. Michel, Márton Megyeri, Maya Schuldiner, Benoît Kornmann, Howard Riezman

## Abstract

Hereditary Sensory and Autonomic Neuropathy (HSAN) type IA and IC are caused by elevated levels of an atypical class of lipid named 1-deoxysphingolipid (DoxSL). How elevated levels of DoxSL perturb the physiology of the cell and how the perturbations lead to HSAN IA/C are largely unknown. In this study, we show that C_26_-1-deoxydihydroceramide (C_26_-DoxDHCer) is highly toxic to the cell, while C_16_- and C_18_-DoxDHCer are less toxic. Genome-wide genetic screens and lipidomics revealed the dynamics of DoxSL accumulation and DoxSL species responsible for the toxicity over the course of DoxSL accumulation. Furthermore, we show that disruption of F-actin organization, alteration of mitochondrial shape, and accumulation of hydrophobic bodies by DoxSL are not sufficient to cause complete cellular failure. However, combined with collapsed ER membrane, these perturbations cause cell death. Thus, we have unraveled key principles of DoxSL cytotoxicity that help to explain the clinical features of HSAN IA/C.

## Introduction

HSAN type IA and IC are inherited diseases that mainly affect the sensory and autonomic functions of the peripheral nervous system. The clinical hallmark of the diseases is loss of pain and temperature sensations in the distal extremities. In some cases, it is accompanied by hypohidrosis (diminished sweating) [1]. In general, the diseases have a late onset varying between the second and the fifth decade of life [1], although cases with a congenital or childhood onset have been reported [2, 3]. The progression of the diseases is usually slow. As the diseases progress, the affected individuals often develop complications, such as ulcerative mutilations; muscle wasting and weakness; reduced motor functions; and spontaneous shooting, burning, and lancinating pains [1, 3, 4], leading to severe physical disabilities.

HSAN IA and IC are caused by autosomal dominant missense mutations in two essential genes, *SPTLC1* and *SPTLC2* [5], respectively. The genes encode the two main subunits of serine palmitoyltransferase (SPT), which is the first enzyme regulating the flux of lipids in the sphingolipid biosynthesis pathway. SPT condenses palmitoyl-CoA with *L*-serine to produce a sphingoid base, 3-ketosphinganine which is rapidly reduced to sphinganine (Sa). Sa can be acylated by ceramide synthase to produce a ceramide, dihydroceramide (DHCer), which can be modified further to generate more complex sphingolipids. Alternatively, Sa can be phosphorylated and then degraded. The latter route constitutes the sphingolipid degradation pathway [6, 7] (**Fig 1A**).

**Fig 1.**
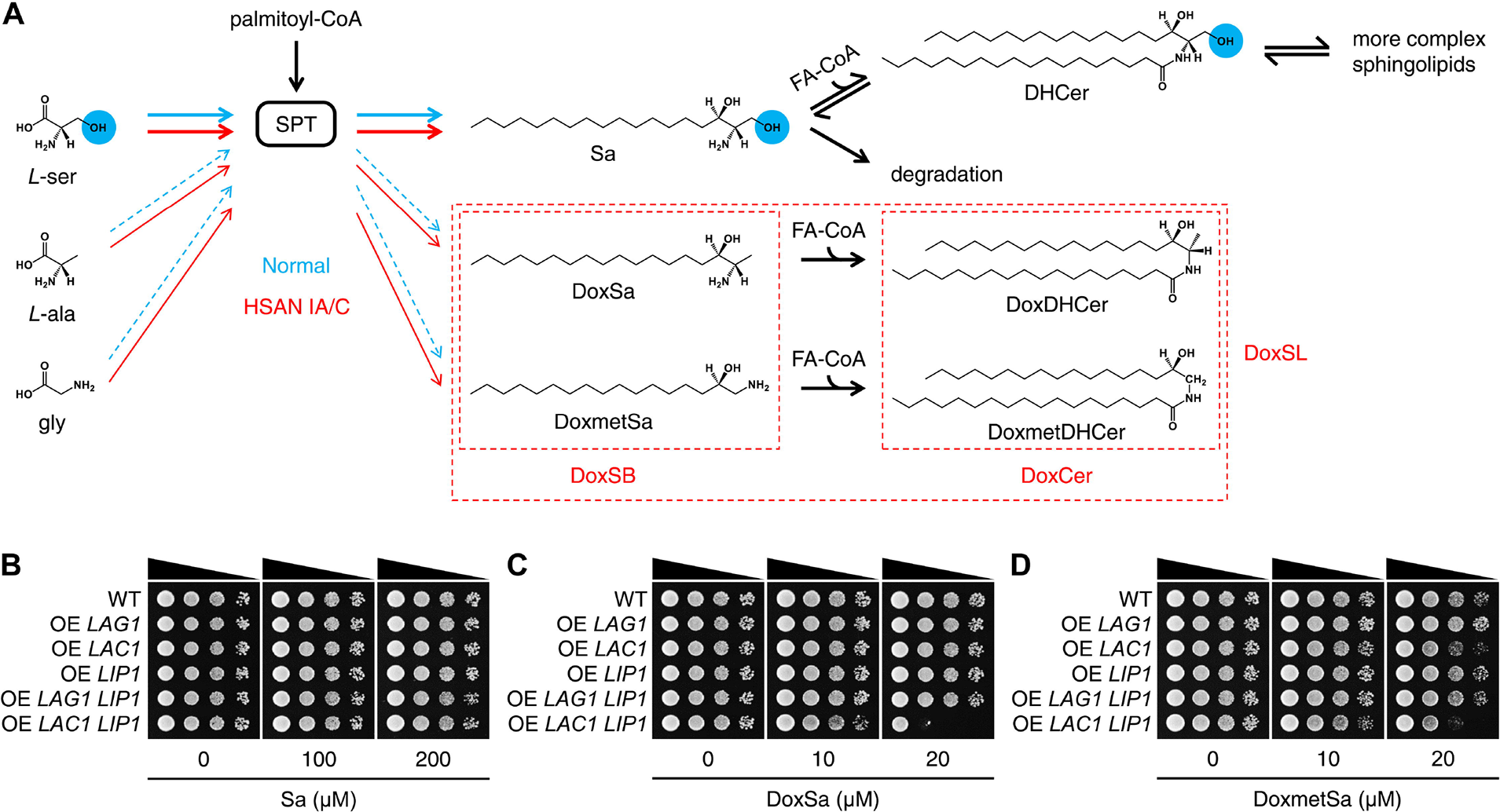
Elevated levels of DoxSL are toxic to yeast. **(A)** Schematic effect of HSAN IA/C mutations on the substrate promiscuity of serine palmitoyltransferase (SPT). The blue circles highlight the hydroxyl group that is missing in 1-deoxysphingolipid (DoxSL). **(B-D)** Effect of sphingoid bases on the growth of the indicated strains evaluated by a spot assay. Sa: sphinganine, DoxSa: 1-deoxysphinganine, DoxmetSa: 1-deoxymethylsphinganine, DoxSB: 1-deoxysphingoid base, FA-CoA: fatty acyl-CoA, DHCer: dihydroceramide, DoxDHCer: 1-deoxydihydroceramide, DoxmetDHCer: 1-deoxymethyldihydroceramide, DoxCer: 1-deoxyceramide.

SPT can also use *L*-alanine or glycine as a substrate, albeit at much lower efficiency, to produce atypical sphingoid bases, 1-deoxysphinganine (DoxSa) or 1-deoxymethylsphinganine (DoxmetSa), respectively. Similar to Sa, DoxSa and DoxmetSa can also be acylated by ceramide synthase to form atypical ceramides, 1-deoxydihydroceramide (DoxDHCer) and 1-deoxymethyldihydroceramide (DoxmetDHCer), respectively. In contrast to the typical sphingolipid, 1-deoxysphingolipid (DoxSL) lacks a hydroxyl group at the first carbon. Since the hydroxyl group is required for the synthesis of complex sphingolipids and the degradation of sphingolipids, DoxSL cannot progress in the pathway and tends to accumulate in the cell [6]. The mutations found in individuals with HSAN IA/C enhance the substrate promiscuity of SPT towards *L*-alanine and glycine [8, 9], leading to increased synthesis and therefore accumulation of DoxSL, leading to HSAN IA/C (**Fig 1A**). How elevated levels of DoxSL perturb the physiology of the cell and how the perturbations lead to disease progression are largely unknown.

Studies in various mammalian cells showed that elevated levels of DoxSL perturb multiple components of the cell, including actin stress fibers [10], mitochondria [11, 12], and lipid droplets [13, 14]. However, to which extent the perturbations affect cell viability is unknown. An inhibitor of ceramide synthase, Fumonisin B1 alleviates the toxic effects of DoxSL [15–17]. In addition, the levels of C_22-24_-1-deoxyceramide (DoxCer) in blood plasma associate with the incidence and severity of neuropathy caused by paclitaxel in cancer chemotherapy [18]. These findings suggest that different species of DoxCer have different degrees of toxicity. Various pathways of DoxSL-induced cell death have been proposed, including ER stress-induced cell death [9, 11], aberrant Ca^2+^ homeostasis-induced apoptosis [12], non-canonical apoptosis [19], and necrosis [16].

In this study, we sought a comprehensive understanding of the cytotoxicity of DoxSL by genome-wide genetic screens and lipidomics. Given that elevated levels of DoxSL are toxic to various mammalian cells, worms [20], fruit flies [21], mice [22], and humans, we hypothesized that elevated levels of DoxSL are toxic to all eukaryotic cells in a conserved manner. Therefore, we used the budding yeast, *Saccharomyces cerevisiae*, (hereafter termed yeast) as a simple eukaryotic model system to reveal the principles of DoxSL toxicity and the key DoxSL-induced perturbations that lead to cell death. By making use of the fatty acyl-CoA specificity of mammalian ceramide synthase, we showed that C_26_-DoxDHCer is more toxic than C_16_- or C_18_-DoxDHCer. Genome-wide genetic screens and lipidomics revealed the dynamics of DoxSL accumulation and DoxSL species responsible for the toxicity over the course of DoxSL accumulation. Furthermore, we showed that DoxSa accumulation leads to depletion of major membrane lipids. By standardizing the conditions of DoxSa treatment, we showed that disruption of F-actin organization, alteration of mitochondrial shape, and accumulation of hydrophobic bodies by DoxSL are not lethal. However, combined with collapsed ER membrane, the perturbations pass the threshold to promote cell death. Thus, we have unraveled key principles of DoxSL cytotoxicity that might explain the clinical features of HSAN IA/C.

## Results

### Elevated levels of DoxSL are toxic to yeast

To test whether elevated levels of 1-deoxysphingolipid (DoxSL) are toxic to yeast, we compared the effect of the typical sphingoid base, sphinganine (Sa) with that of 1-deoxysphingoid base (DoxSB) on cell growth by a spot assay. To ensure that the formation of 1-deoxyceramide (DoxCer) from supplemented DoxSB is not limited by the endogenous ceramide synthase, we overexpressed the enzyme by integrating an additional copy of its gene with the constitutively active strong promoter of glyceraldehyde-3-phosphate dehydrogenase isozyme 3 (*TDH3*) into the yeast genome. Yeast ceramide synthase may be composed of a Lag1p or Lac1p homodimer or a Lag1p-Lac1p heterodimer. Each of these dimers contains at least two copies of Lip1p, which is an essential protein required for the activity of yeast ceramide synthase [23]. Therefore, we also overexpressed *LIP1* in the same way.

The assay showed that Sa did not inhibit cell growth regardless of ceramide synthase overexpression up to 200 μM (**Fig 1B**). In contrast to Sa, 1-deoxysphinganine (DoxSa) was sufficient to strongly inhibit the growth of the strain overexpressing *LAC1 LIP1* (**Fig 1C**) at 20 μM. Similarly, 1-deoxymethylsphinganine (DoxmetSa) was also sufficient to inhibit the growth of the same strain at the same concentration, albeit to a lesser degree (**Fig 1D**). This result demonstrates that DoxSB is much more toxic than the typical sphingoid base and suggests that elevated levels of DoxCer are also toxic to yeast.

At the same concentration, however, DoxSa or DoxmetSa did not inhibit the growth of the strain overexpressing *LAG1 LIP1*. It is possible that Lag1p is less abundant than Lac1p because the expression level of *LAC1* is normally higher than that of *LAG1* [24] or that Lag1p is less efficient than Lac1p in converting DoxSB to DoxCer. The latter explanation is consistent with the recent finding that Lag1p prefers phytosphingosine to Sa as a substrate (Megyeri et al., 2019, *in press*), and therefore is expected to have reduced activity towards DoxSa or DoxmetSa.

### C_26_-DoxDHCer is more toxic than C_16_- or C_18_-DoxDHCer

To evaluate the toxicity of DoxCer species differing in the length of their acyl chain, we made use of the fatty acyl-CoA specificity of mammalian ceramide synthase. There are six ceramide synthases (CerS1-6) in mammals. Each enzyme uses specific fatty acyl-CoA species to produce corresponding ceramide species [25] (**Fig 2A**). We expressed individual mammalian ceramide synthases in yeast to confer on yeast an ability to convert supplemented DoxSB to different DoxCer species. The enzymes were expressed by integrating their genes with the *TDH3* promoter into the yeast genome. Each of the expressed enzymes was able to support growth when the genes of yeast ceramide synthase were deleted (*Δlag1 Δlac1*), demonstrating that they all are functional in yeast (**S1A Fig**). We then subjected the strains expressing mammalian ceramide synthases in addition to the endogenous ones to a spot assay.

**Fig 2.**
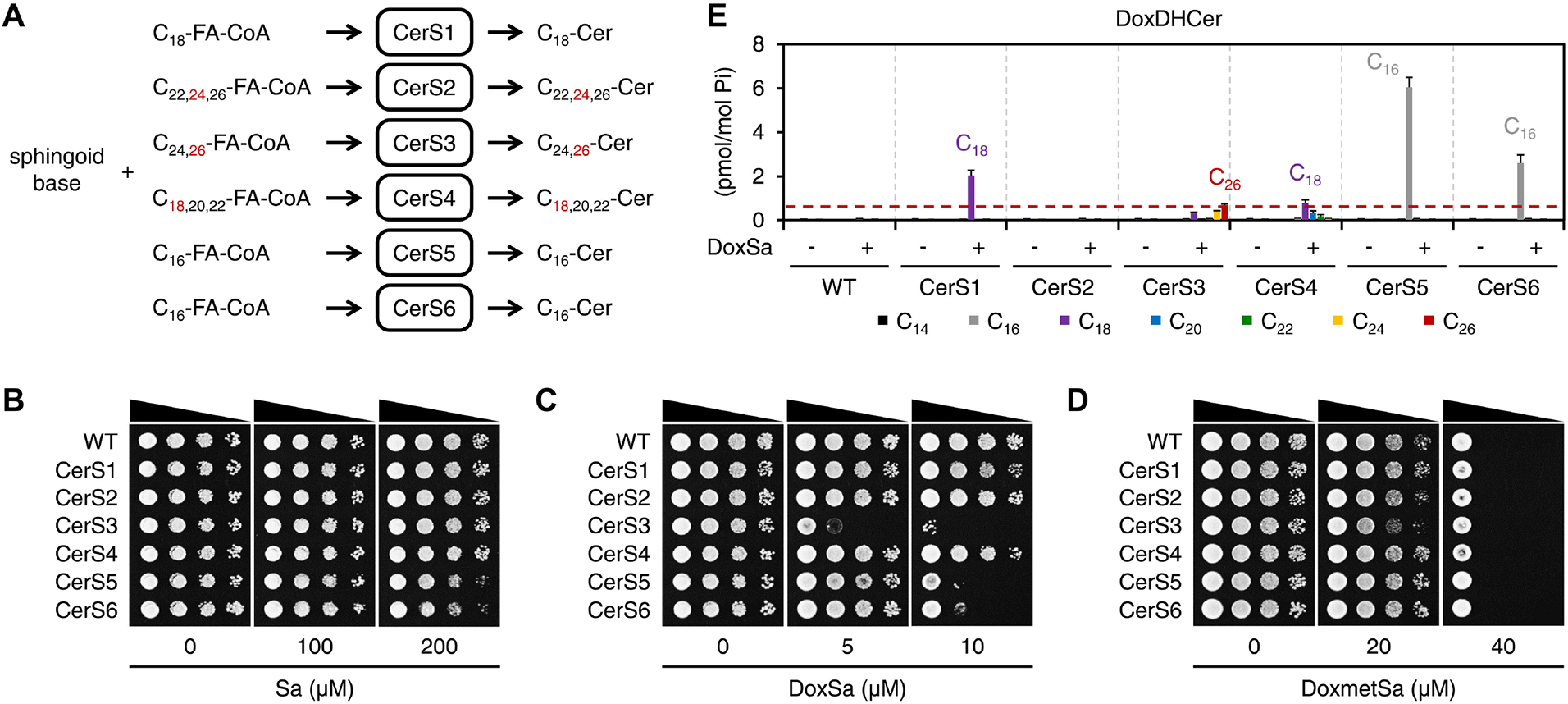
C_26_-DoxDHCer is more toxic than C_16_- or C_18_-DoxDHCer. **(A)** Schematic representation of the fatty acyl-CoA specificity of mammalian ceramide synthase. The red numbers indicate the major species of fatty acyl-CoA and ceramide (Cer). **(B-D)** Effect of sphingoid bases on the growth of the indicated strains evaluated by a spot assay. **(E)** Levels of DoxDHCer species in the indicated strains without or with a non-toxic DoxSa treatment (2 μM of DoxSa, 20 million cells/ml, 1.5 h) determined by mass spectrometry (MS). The red dashed line indicates the level of C_26_-DoxDHCer in the CerS3 strain following the treatment.

The assay showed that Sa at 200 μM did not inhibit the growth of WT and the strains expressing CerS1, 2, 3, or 4, and mildly inhibited the growth of the strains expressing CerS5 or 6 (**Fig 2B**). In contrast to Sa, DoxSa at 5 μM was sufficient to strongly inhibit the growth of the strain expressing CerS3. At 10 μM, DoxSa strongly inhibited the growth of the strains expressing CerS3, 5 or 6, but did not inhibit the growth of WT nor the strains expressing CerS1, 2, or 4 (**Fig 2C**). This result suggests that the toxicity of 1-deoxydihydroceramide (DoxDHCer) depends on the length of its acyl chain. In contrast to DoxSa, DoxmetSa at 40 μM strongly inhibited the growth of WT and all strains to the same degree (**Fig 2D**). This result suggests either that mammalian ceramide synthase expressed in yeast cannot convert DoxmetSa to 1-deoxymethyldihydroceramide (DoxmetDHCer) or that DoxmetSa is toxic on its own. Therefore, we cannot evaluate the toxicity of different DoxmetDHCer species.

To determine the species and the levels of accumulated DoxDHCer in cells for a given concentration of DoxSa, we treated the strains with DoxSa in a liquid medium at a non-toxic condition (low DoxSa concentration, high cell density, one yeast generation time) and then measured the levels of sphingolipid in the cells by mass spectrometry (MS). A non-toxic condition was chosen to ensure that the cells were not saturated with DoxDHCer, therefore allowing comparison between the levels of accumulated DoxDHCer in different strains. The MS analysis showed that DoxSa had a minor effect on the levels of typical sphingolipid (**S1B-H Fig**), demonstrating that the treatment condition is not toxic to the cells. The analysis also showed that following the DoxSa treatment, the strain expressing CerS3 mainly accumulated C_26_-DoxDHCer. Interestingly, the level of C_26_-DoxDHCer in the strain was much higher than that in WT, indicating that the mammalian CerS3, which makes the same DHCer species as the yeast ceramide synthase, is more efficient in producing C_26_-DoxDHCer than the yeast enzyme. Moreover, the level of C_26_-DoxDHCer in the strain was lower than those of C_16_-DoxDHCer in the strains expressing CerS5 or 6 and those of C_18_-DoxDHCer in the strains expressing CerS1 or 4 (**Fig 2E**). Nevertheless, the strain expressing CerS3 was the most sensitive strain to DoxSa (**Fig 2C**). These results strongly indicate that C_26_-DoxDHCer is more toxic than C_16_- or C_18_-DoxDHCer.

### Genome-wide genetic screen of knockout and hypomorphic (DAmP) mutants

To gain insights into the mechanism of action (MoA) of DoxSL, we performed a genome-wide genetic screen for gene products whose absence or deficiency render the cell hypersensitive or resistant to DoxSL. We decided to focus on DoxSa and DoxDHCer for two reasons. First, we have more information about the effect of acyl chain length on the toxicity of DoxDHCer than that of DoxmetDHCer. Second, we could minimize the amount of DoxSa consumed for the screen by using the strain expressing CerS3, which is the most sensitive strain to DoxSa, as a background strain. The screen was performed by first introducing CerS3 into every single yeast strain in the knockout or hypomorphic (DAmP) allele library using automated approaches [26, 27]. Then, the resulting library, in which each strain expressed CerS3 and had one mutant allele, was replica plated onto an agar medium supplemented with different concentrations of DoxSa. Two concentrations were chosen, such that hypersensitive or resistant mutants were revealed prominently at the lower or the higher concentration, respectively. Colony size was scored as a measure of fitness of the mutants (**Fig 3A**).

**Fig 3.**
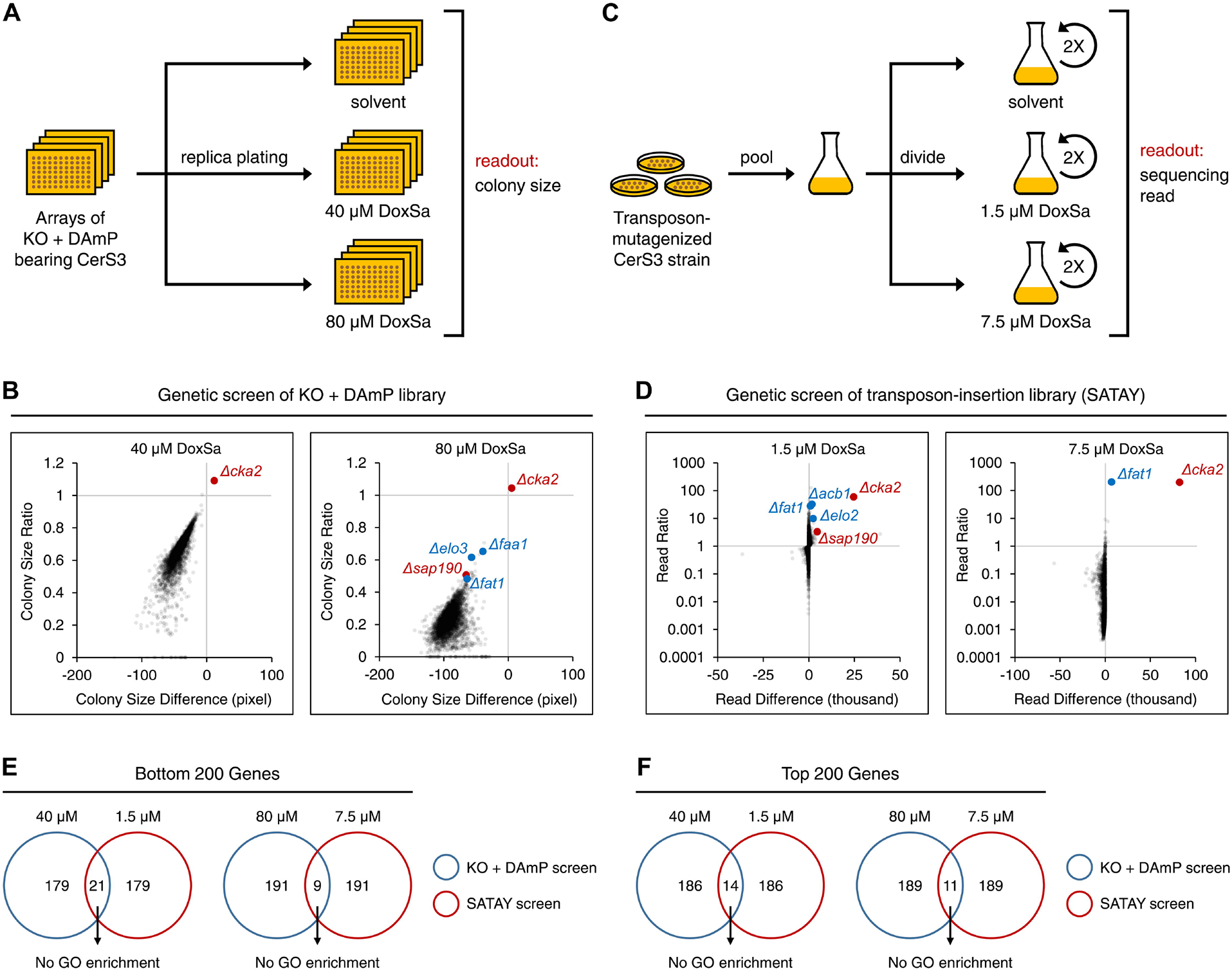
Genome-wide genetic screens to reveal the mechanism of action of DoxSL. **(A)** Schematic workflow of genome-wide genetic screen of knockout and hypomorphic (DAmP) mutants. **(B)** Changes in colony size of the mutants following a DoxSa treatment at the indicated concentrations. **(C)** Schematic workflow of genome-wide genetic screen of transposon-insertion mutants (SATAY). **(D)** Changes in sequencing read of the mutants following a DoxSa treatment at the indicated concentrations. **(E and F)** Proportions of genes that were in the bottom **(E)** or top **(F)** 200 of the fitness ranks and that were revealed by the two genetic screens. There is no gene ontology (GO) enrichment in each set of genes.

The library covered 79.5% of non-essential genes and 68.8% of essential genes (**S2A Fig**). Quality control assessments revealed that the extents of biases and technical variabilities in the screen were below our tolerance limits (**S2B-F Fig**). The screen revealed that deletion of genes that are required for efficient import of sphingoid base (*Δfaa1*) or synthesis of very long-chain fatty acyl-CoA (*Δelo3, Δfat1*) rendered the cell resistant to DoxSa (**Fig 3B**). Since efficient synthesis of C_26_-DoxDHCer from DoxSa requires the two processes, this finding validates that changes in fitness of the mutants in the screen were mainly due to the high toxicity of C_26_-DoxDHCer.

Gene ontology (GO) enrichment analysis of resistant genes (defined as genes whose Z-score of colony size ratio > 3, Z-score of solvent-treated colony size > −3.5, CV of colony size ratio < the 50% percentile) using the GOrilla tool [28, 29] showed that there is no enrichment of particular molecular functions, cellular components, or biological processes. Moreover, physical and genetic-interaction density analysis [30] was unable to reveal groups of functionally related hypersensitive or resistant genes. In addition, examination of individual hypersensitive or resistant genes failed to reveal a clear hypothesis of the MoA of DoxSL. During the examination of individual genes, we found that the identities of about 10% of the mutants in the library could not be confirmed by colony PCR. Furthermore, several freshly generated mutants failed to show the same responses to DoxSa as those indicated by the result of the screen. These findings raised a concern that the mutants in the library had accumulated too many suppressors, thus preventing us from obtaining insights into the MoA of DoxSL.

### Genome-wide genetic screen of transposon-insertion mutants (SATAY)

To ensure that we did not miss key gene products involved in the MoA of DoxSL, we decided to perform another genome-wide genetic screen with an independent method in which the library is freshly-generated. The method is termed SAturated Transposon Analysis in Yeast (SATAY) [31]. The screen was performed by first inducing random insertion of a transposon from a plasmid into the genome of the CerS3 strain. The mutagenesis was performed in a large number of cells to generate a saturated transposon-insertion library. Then, the library was divided and subjected to a two-round treatment with DoxSa at different concentrations. Two concentrations were chosen for the same purpose as that in the screen of knockout and DAmP mutants. The number of deep sequencing reads was used as a measure of fitness of the mutants (**Fig 3C**).

The transposon insertion mutated 91.5% of non-essential genes and 52.5% of essential genes (**S3A Fig**). The number of transposon insertion sites of most genes remained unchanged following DoxSa treatment at the lower concentration (**S3B Fig**), suggesting that the complexity of the library was maintained during the screening procedure and that technical variabilities had little impact on the result of the screen. The screen revealed that disruption of genes that are required for synthesis of very long-chain fatty acyl-CoA and its presentation to the ceramide synthase (*Δfat1, Δacb1, Δelo2*) rendered the cell resistant to DoxSa (**Fig 3D**). This finding validates that changes in fitness of the mutants in the screen were mainly due to the high toxicity of C_26_-DoxDHCer.

We performed various data analyses ranging from enrichment analyses to examination of individual transposon insertion sites. Furthermore, we scrutinized genes that were in the bottom or top 200 of the fitness ranks and that were revealed by the two genetic screens (**Fig 3E and 3F**). However, once again, we were unable to formulate a clear hypothesis of the MoA of DoxSL. To allow independent analyses of the datasets by others, all primary data are published here (**S1 and S2 Tab**). It is possible that the MoA cannot be revealed by genetic screens in which gene products are deleted, disrupted, or reduced. Therefore, we performed a multicopy suppressor screen for gene products whose overproduction alleviates the toxicity of DoxSL. In this screen, the CerS3 strain was transformed with a pool of plasmids, each of which carried 4-5 genes and the 2-micron sequence that maintains a high copy number of the plasmid in the cell. Then, the transformed cells were subjected to a three-round treatment with DoxSa at different concentrations. We found that cultures treated with high concentrations of DoxSa did not gain noticeable density after the second and the third round of treatment (data not shown). Considering that the plasmid library covers 97.2% of the yeast genome with 5.4-fold depth of coverage [32] and that the transformed cells covered the plasmid library with 15.9-fold depth of coverage, this finding suggests that there is no single gene whose overexpression is capable of alleviating the toxicity of DoxSL.

Given the results of the three genetic screens, we hypothesized two possibilities of the MoA of DoxSL. (1) DoxSL inhibits an essential multisubunit protein complex and the inhibition cannot be suppressed by deletion or overproduction of another protein. (2) DoxSL inhibits multiple proteins and pathways, such that perturbing any single one does not reveal the full MoA. This inhibition could be achieved directly or indirectly, such as by affecting the physical properties of cellular membranes.

### The dynamics of DoxSL species responsible for the toxicity over the course of DoxSL accumulation

To gain more insights into the toxicity of DoxSL, we subjected the deletion mutants of the resistant genes required for efficient import of sphingoid base or synthesis of very long-chain fatty acyl-CoA, along with those of two resistant genes, *CKA2* (encoding alpha catalytic subunit of casein kinase 2) and *SAP190* (encoding a component of type 2A-related serine-threonine phosphatase SIT4) to a spot assay and to MS analysis following the non-toxic DoxSa treatment.

The assay showed that all mutants were more resistant, to different degrees, to DoxSa at 5 μM than the parent strain (the CerS3 strain) (**Fig 4A**). The MS analysis showed that all mutants accumulated less C_26_-DoxDHCer than the CerS3 strain following the DoxSa treatment (**Fig 4B**). It also showed that C_26_-DoxDHCer was the only DoxDHCer species whose accumulation levels in all mutants were lower than that in the CerS3 strain (**S4A-S4F Fig**). Therefore, the degrees of resistance of the mutants could be attributed to their ability to suppress the accumulation of C_26_-DoxDHCer. However, the degree of resistance and the ability to suppress the accumulation of C_26_-DoxDHCer of the mutants did not correlate perfectly, suggesting that C_26_-DoxDHCer was not the only toxic DoxSL species. This became more apparent at a higher concentration of DoxSa. The degrees of resistance of the mutants at 10 μM of DoxSa (**Fig 4A**) could not be explained by their ability to suppress the accumulation of C_26_-DoxDHCer (**Fig 4B**). Instead, they could be better explained by their ability to suppress the accumulation of all DoxDHCer species, albeit imperfectly (**Fig 4C**).

**Fig 4.**
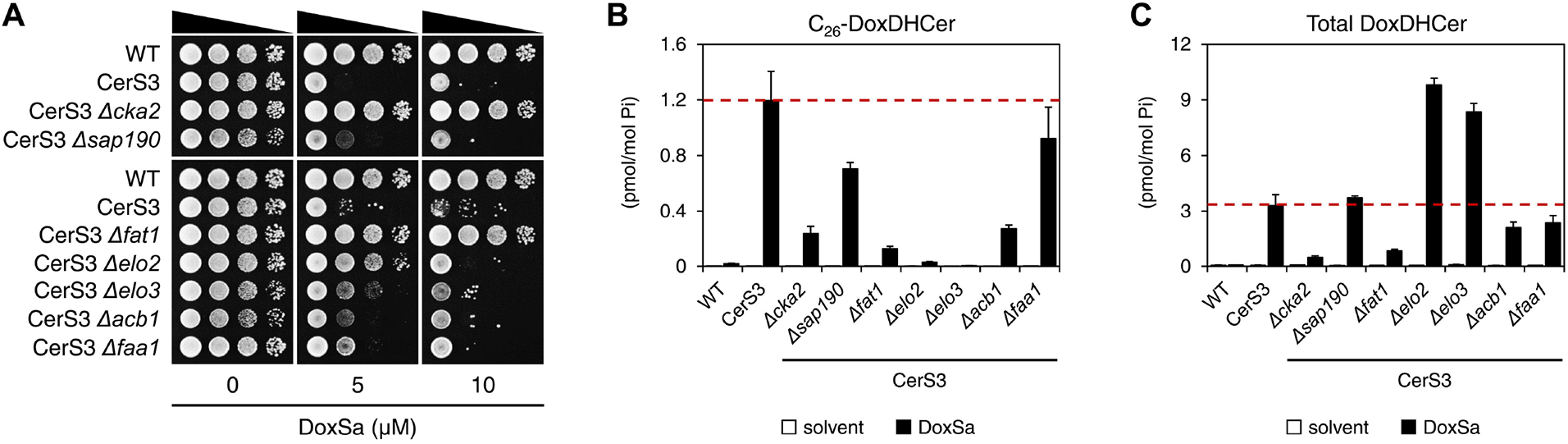
The dynamics of DoxSL species responsible for the toxicity over the course of DoxSL accumulation. **(A)** Effect of DoxSa on the growth of the indicated strains evaluated by a spot assay. **(B and C)** Levels of C_26_-DoxDHCer **(B)** and total DoxDHCer **(C)** in the indicated strains without or with a non-toxic DoxSa treatment (2 μM of DoxSa, 20 million cells/ml, 1.5 h) determined by MS. The red dashed lines indicate the levels of C_26_-DoxDHCer and total DoxDHCer in the CerS3 strain following the treatment.

The results indicate that the toxicity of DoxSL is the summed toxicity of DoxSa and individual DoxDHCer species. The results also show that the DoxSL species responsible for the toxicity varies depending on the amount of DoxSL that accumulates. When DoxSL is present at low levels, its toxic effects are mainly caused by the most toxic DoxDHCer species (C_26_-DoxDHCer.) When DoxSL is present at high levels, its toxic effects cannot be attributed to one main species as the toxic effects of DoxSa and the other DoxDHCer species become more evident.

### Standardizing conditions of DoxSa treatment for phenotypic characterization

To gain more insights into the toxicity of DoxSL, we characterized the phenotypes of DoxSL toxicity. To this end, we first standardized conditions of DoxSa treatment by testing the effect of various concentrations of DoxSa, durations of DoxSa treatment, and cell densities on cell viability. We aimed to obtain a range of DoxSa concentrations that gives a wide range of cell viabilities after one yeast generation time (1.5 h). One generation time was chosen to maximize the effects of DoxSL while minimizing the compounding effects of inter-generation events and the depletion of DoxSa from the medium. We treated cells at low cell densities to maximize their sensitivity to DoxSL. Following a treatment, cells were washed and spotted on an agar medium to assay their ability to recover. We found that treating WT and the CerS3 strain with DoxSa at concentrations of 0, 2, 4, and 6 μM, for 1.5 h, at a cell density of 1 million cells/ml gave a wide range of cell viabilities (**Fig 5A**). Therefore, we used these conditions of DoxSa treatment for the phenotypic characterization.

**Fig 5.**
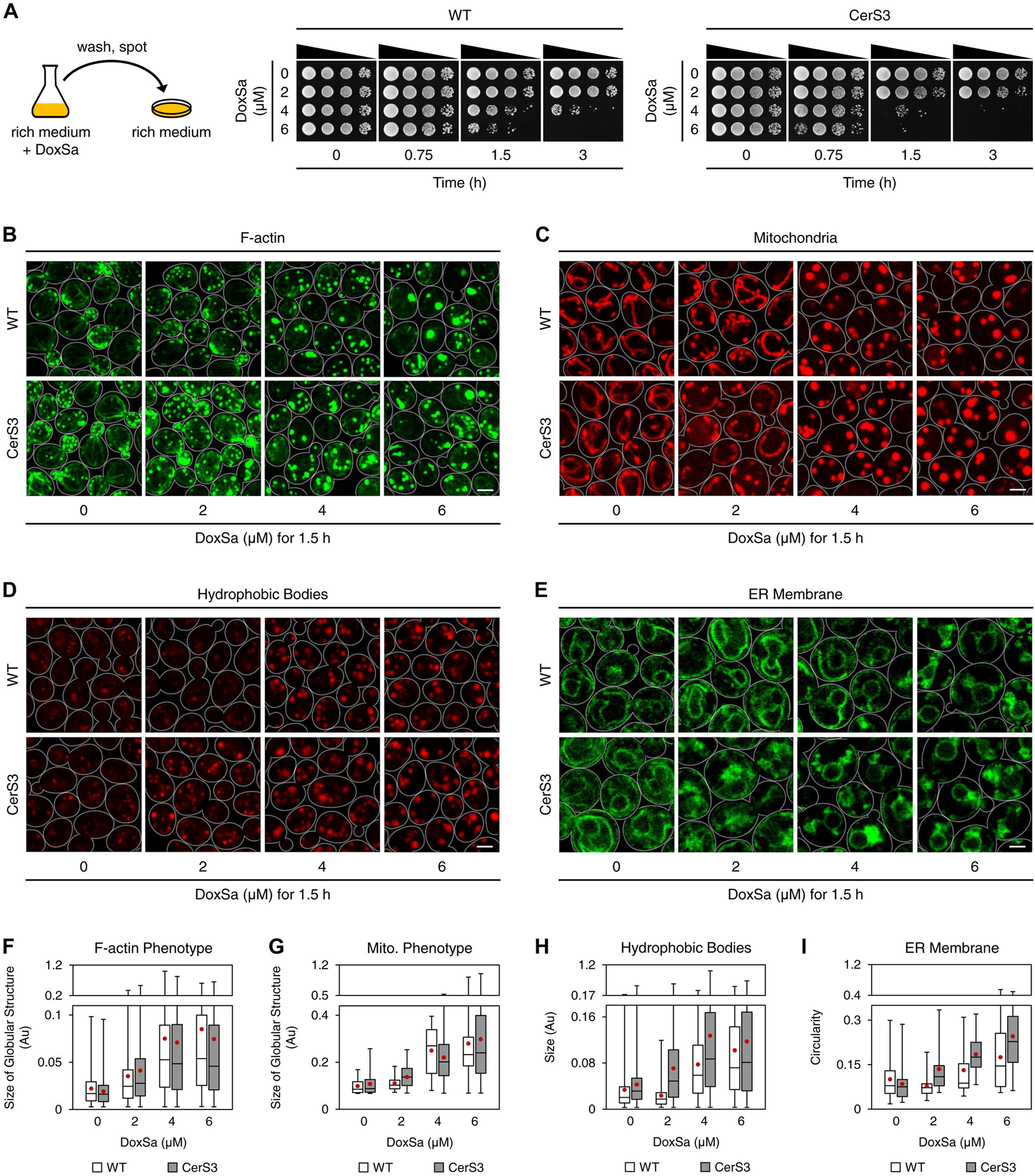
Cellular phenotypes of DoxSL toxicity. **(A)** Ability of WT and the CerS3 strain to recover from the indicated DoxSa treatments. **(B-E)** Cellular phenotypes of DoxSL toxicity. Organization of F-actin **(B)**, shape of mitochondria **(C)**, hydrophobic bodies **(D)**, and the ER membrane **(E)** in WT and the CerS3 strain following standardized DoxSa treatments (indicated concentrations of DoxSa, 1 million cells/ml, 1.5 h). F-actin was stained with phalloidin-Atto488. Mitochondria were visualized with Mdh1p-mCherry. Hydrophobic bodies were stained with Nile Red. The ER membrane was visualized with Sec61p-GFP. Each image is a maximum intensity projection of a 2-μm slice. The scale bars are 2 μm. **(F-I)** Size of globular structures stained with phalloidin-Atto488 **(F)**, size of globular structures highlighted by Mdh1p-mCherry **(G)**, size of hydrophobic bodies **(H)**, or circularity (a value of 1.0 indicates a perfect circle) of the ER membrane **(I)** in WT and the CerS3 strain following the standardized DoxSa treatments. Black bars indicate medians, boxes indicate quartiles, whiskers indicate extreme values, and red dots indicate means.

### Disruption of F-actin organization is not sufficient to cause cell death

DoxSa treatment was shown to reduce the presence of actin stress fibers in a mammalian cell line [10]. To test whether DoxSa treatment also disrupts F-actin organization in yeast, we performed confocal microscopy using phalloidin-Atto488 as a stain for F-actin. The microscopy showed that 4 μM of DoxSa for 1.5 h was sufficient to depolarize actin patches and markedly reduce the presence of actin cables and actin patches in WT. Moreover, it induced the formation of multiple micron-sized round bodies that could be stained with phalloidin-Atto488. The microscopy also showed that 2 μM of DoxSa for 1.5 h gave the same phenotype in the CerS3 strain, albeit to a lesser degree (**Fig 5B and 5F**). This result shows that DoxSa treatment disrupts F-actin organization in yeast. However, 4 μM of DoxSa for 1.5 h was not sufficient to stop the growth of WT (**Fig 5A**), demonstrating that the phenotype is not sufficient to cause cell death.

### Alteration of mitochondrial shape is not sufficient to cause cell death

DoxSa treatment was shown to induce mitochondrial fragmentation, induce the loss of mitochondrial cristae, reduce respiration, and reduce ATP production in a mammalian cell line [11]. To test whether DoxSa treatment also perturbs mitochondria in yeast, we performed confocal microscopy of strains expressing mCherry tagged-mitochondrial malate dehydrogenase (*MDH1-mCherry*). The microscopy showed that 4 μM of DoxSa for 1.5 h was sufficient to fragment mitochondria and alter the shape of mitochondria from tubular to spherical without affecting their inheritance from the mother cell to the bud in WT. The microscopy also showed that 2 μM of DoxSa for 1.5 h gave the same phenotype in the CerS3 strain, albeit to a lesser degree (**Fig 5C and 5G**). This result shows that DoxSa treatment perturbs mitochondria in yeast. However, 4 μM of DoxSa for 1.5 h was not sufficient to stop the growth of WT (**Fig 5A**), demonstrating that the phenotype is not sufficient to cause cell death. Although the mitochondrial phenotype was dramatic, depletion of oxygen, non-fermentable carbon sources, or deletion of mitochondrial DNA did not modulate the sensitivity of WT and the CerS3 strain to DoxSa (**S5A-S5F Fig**). These results suggest that the phenotype is independent of mitochondrial respiration.

### Accumulation of hydrophobic bodies is not sufficient to cause cell death

Mammalian cells lacking *de novo* biosynthesis of *L*-serine and HSAN IA patient-derived lymphoblasts were shown to have more lipid droplets than normal cells [13, 14]. To test whether DoxSa treatment also enhances the formation of lipid droplets in yeast, we performed confocal microscopy using a hydrophobic dye, Nile Red. The microscopy showed that 4 μM of DoxSa for 1.5 h was sufficient to markedly induce the formation of Nile Red-stained structures whose sizes, shapes, subcellular locations, and sharpness of the boundaries are distinct from those of lipid droplets (**Fig 5D**). The structures could also be observed in a quadruple mutant lacking the abilities to synthesize triglycerides and sterol esters (*Δare1 Δare2 Δdga1 Δlro1*) following DoxSa treatment (**S6A and S6B Fig**), demonstrating that the formation of the structures is independent of neutral lipids and that they are not the canonical lipid droplets. Therefore, we call the structures “hydrophobic bodies”. The microscopy also showed that 2 μM of DoxSa for 1.5 h gave the same phenotype in the CerS3 strain, albeit to a lesser degree (**Fig 5D and 5H**). These results show that DoxSa treatment induces the formation of hydrophobic bodies in yeast. However, 4 μM of DoxSa for 1.5 h was not sufficient to stop the growth of WT (**Fig 5A**), demonstrating that the phenotype is not sufficient to cause cell death.

### Collapsed ER membrane coincides with cell death

Since the conversion of DoxSa to DoxDHCer presumably occurs at the ER membrane, where ceramide synthase resides [23, 33], we tested the effect of DoxSa treatment on the ER membrane by confocal microscopy of strains expressing GFP tagged-translocon (*SEC61-GFP*). The microscopy showed that 6 μM of DoxSa for 1.5 h was required to markedly collapse the cortical ER in WT. The microscopy also showed that 4 μM of DoxSa for 1.5 h was sufficient to induce more severe collapsed cortical ER in the CerS3 strain (**Fig 5E and 5I**). This result shows that DoxSa treatment causes the ER membrane to collapse in yeast. In contrast to the other phenotypes, marked collapse of the cortical ER concurred with marked reduction of cell viability (**Fig 5A and 5E**), demonstrating that collapsed ER membrane coincides with cell death.

We also tested the effects of DoxSa treatment on the nuclear envelope, the vacuolar membrane, the peroxisomal membrane, and the plasma membrane by confocal microscopy. However, we could not observe apparent morphological changes of the membranes (data not shown), indicating that different cellular membranes are affected by elevated levels of DoxSL to different degrees.

### DoxSa accumulation leads to depletion of major membrane lipids

To gain more insights into the toxicity of DoxSL, we performed lipidomics of WT and the CerS3 strain following the same DoxSa treatments as those for the phenotypic characterization by MS. The lipidomics showed that increased concentrations of DoxSa were accompanied by decreased levels of major membrane lipids in both WT and the CerS3 strain. The lipids were five classes of glycerophospholipid (PC, PI, PS, PE, and CL) (**Fig 6A-6E**) and two classes of complex sphingolipid (IPC and MIPC) (**Fig 6G and 6H**). In contrast, increased concentrations of DoxSa were accompanied by increased levels of a sphingolipid intermediate, ceramide (**Fig 6F**). Compared to those of other lipid classes, the levels of a class of complex sphingolipid, M(IP)_2_C were the least affected by increased concentrations of DoxSa (**Fig 6I**). Given that the degrees of depletion of the major membrane lipids in WT were comparable to those in the CerS3 strain, this result suggests that DoxSa, not DoxDHCer, accumulation leads to depletion of major membrane lipids.

**Fig 6.**
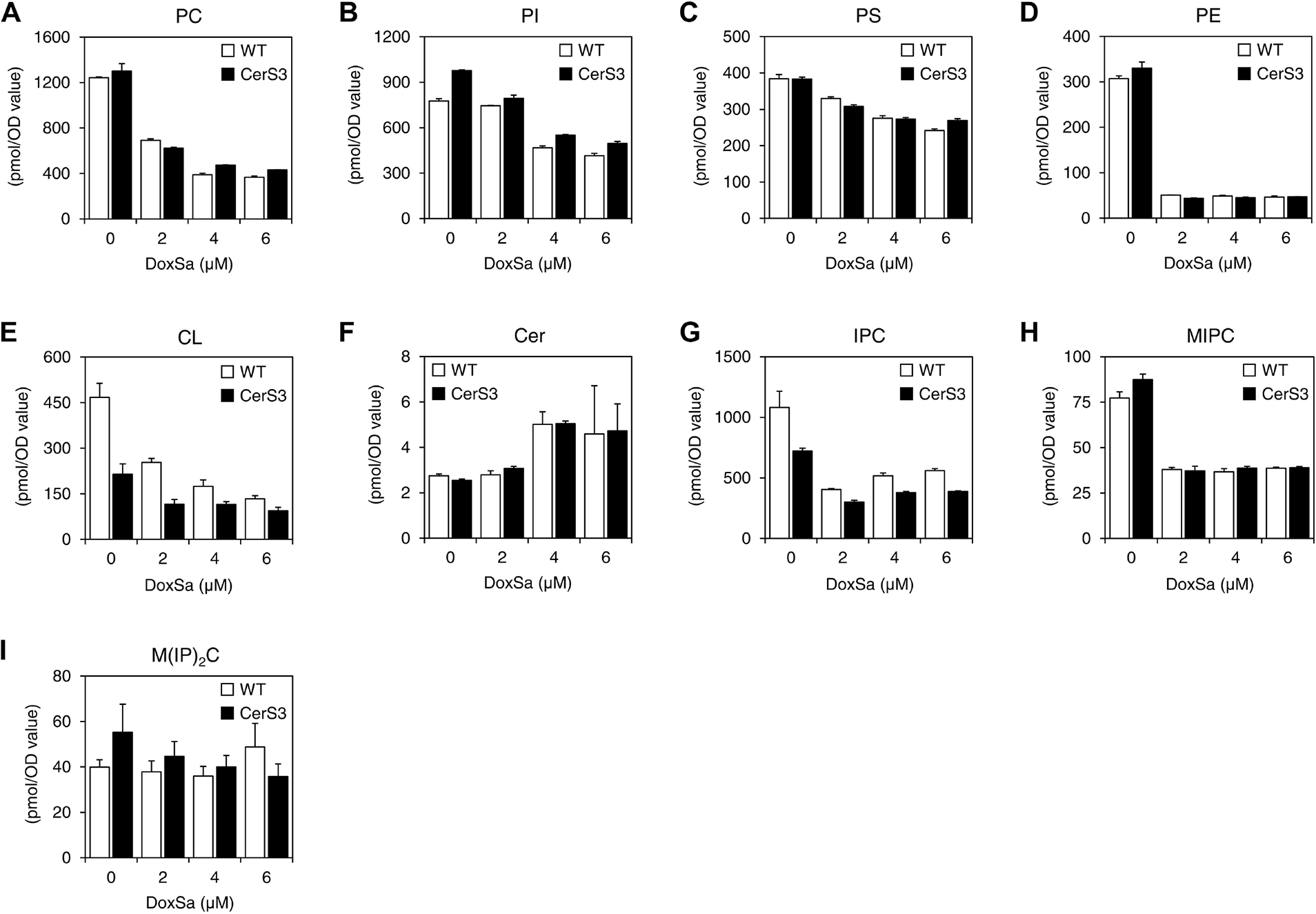
DoxSa accumulation leads to depletion of major membrane lipids. Levels of different categories of phospholipid **(A-E)** and sphingolipid **(F-I)** in WT and the CerS3 strain following the standardized DoxSa treatments (indicated concentrations of DoxSa, 1 million cells/ml, 1.5 h) determined by MS. PC: phosphatidylcholine, PI: phosphatidylinositol, PS: phosphatidylserine, PE: phosphatidylethanolamine, CL: cardiolipin, Cer: ceramide, IPC: inositol-phosphoceramide, MIPC: mannose-inositol-phosphoceramide, M(IP)_2_C: mannose-(inositol-P)_2_-ceramide.

### The dynamics of DoxSL accumulation: the rate of DoxSa accumulation determines the level of DoxDHCer accumulation

The lipidomics also showed that increased concentrations of DoxSa were accompanied by increased levels of C_26_-DoxDHCer and total DoxDHCer in the CerS3 strain and, to lesser degrees, in WT. In the CerS3 strain, we uncovered an unexpected phenomenon in which 2 μM of DoxSa resulted in higher levels of C_26_-DoxDHCer and total DoxDHCer than 4 or 6 μM of DoxSa (**Fig 7A and 7B**). A time-course lipid analysis by MS following a DoxSa treatment at 6 μM for up to 3 h showed that the longer the DoxSa treatment, the higher the levels of C_26_-DoxDHCer and total DoxDHCer (**Fig 7C and 7D**), demonstrating that the phenomenon is not due to post-cell death events. In addition, deletion of ceramidase (*Δypc1 Δydc1*) did not modulate the sensitivity of the CerS3 strain to DoxSa (**Fig 7E**), demonstrating that the phenomenon is ceramidase-independent. The phenomenon suggests the dynamics of DoxSL accumulation – that is to say, the rate of DoxSa accumulation determines the level of DoxDHCer accumulation. When the rates of DoxSa accumulation are high, the toxic effects of DoxSa rapidly manifest, thereby hindering further synthesis and accumulation of DoxDHCer. When the rates of DoxSa accumulation are low, the toxic effects of DoxSa never or slowly manifest, thereby allowing continuous or prolonged synthesis and accumulation of DoxDHCer. Since CerS3 allows efficient conversion of DoxSa to DoxDHCer, the phenomenon is more apparent in the CerS3 strain.

**Fig 7.**
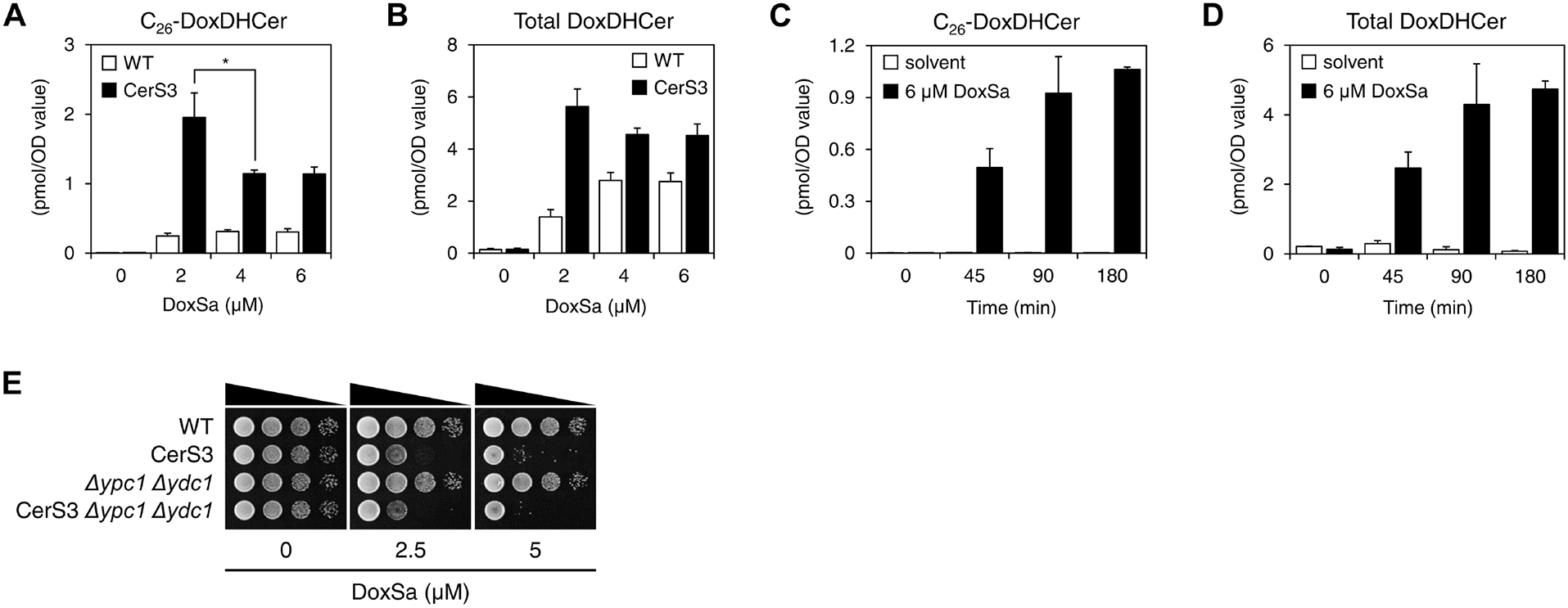
The rate of DoxSa accumulation determines the level of DoxDHCer accumulation. Levels of C_26_-DoxDHCer **(A)** and total DoxDHCer **(B)** in WT and the CerS3 strain following the standardized DoxSa treatments (indicated concentrations of DoxSa, 1 million cells/ml, 1.5 h) determined by MS. **(C and D)** Levels of C_26_-DoxDHCer **(C)** and total DoxDHCer **(D)** in the CerS3 strain monitored by a time-course lipid analysis by MS following a DoxSa treatment at 6 μM for up to 3 h. **(E)** Effect of DoxSa on the growth of the indicated strains evaluated by a spot assay. Deletion of ceramidase (*Δypc1 Δydc1*) did not modulate the sensitivity of the CerS3 strain to DoxSa, indicating that the accumulation of DoxDHCer is independent of ceramidase.

### Model of the cytotoxicity of DoxSL

Since the phenotypes in the CerS3 strain were more severe than those in WT following a given DoxSa treatment (**Fig 5B-5I**), the phenotypes could not be fully attributed to elevated levels of DoxSa and depletion of major membrane lipids (**Fig 6A-6I**). Also, since the levels of accumulated DoxDHCer did not fully correlate with the severity of the phenotypes (**Fig 5B-5I, Fig 7A and 7B**), the phenotypes could not be fully attributed to elevated levels of DoxDHCer either. Therefore, the phenotypes must be caused by elevated levels of both DoxSa and DoxDHCer. Furthermore, deletion of the ER-to-plasma membrane tethers was shown to cause a similarly collapsed ER membrane phenotype without causing cell death in yeast [34]. Therefore, it is likely that in this case as well that the collapsed ER membrane is not sufficient to cause cell death, although it coincides with cell death.

Collectively, our findings suggest the following model of the cytotoxicity of DoxSL. Elevated levels of both DoxSa and DoxDHCer disrupt the organization of F-actin, alter the shape of mitochondria, induce the formation of hydrophobic bodies, and cause the ER membrane to collapse. The first three perturbations reduce cell viability, but they are not sufficient to cause complete cellular failure. The combination of all four perturbations leads to cell death (**Figure 8**). Considering that (1) DoxSa and individual DoxDHCer species might have different mechanisms of action and that (2) the perturbations are expected to negatively affect multiple proteins and pathways, this model is more consistent with our second hypothesis formulated from the results of the three genetic screens.

**Fig 8.**
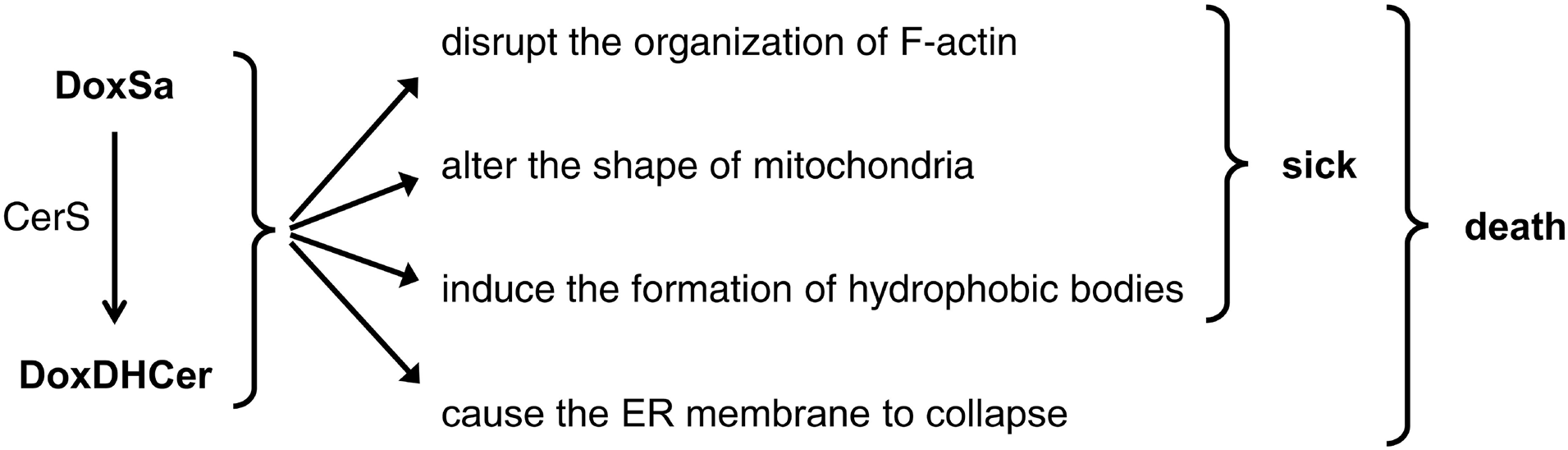
Model of the cytotoxicity of DoxSL. Elevated levels of both DoxSa and DoxDHCer disrupt the organization of F-actin, alter the shape of mitochondria, induce the formation of hydrophobic bodies, and cause the ER membrane to collapse. The first three perturbations reduce cell viability, but they are not sufficient to cause complete cellular failure. The combination of all four perturbations leads to cell death. This model is consistent with the hypothesis that DoxSL inhibits multiple proteins and pathways. This inhibition could be achieved directly or indirectly, such as by affecting the physical properties of cellular membranes.

## Discussion

In this study, we took comprehensive approaches to the cytotoxicity of DoxSL. Gradual accumulation of DoxSL underlies late-onset slowly-progressing neurological diseases HSAN IA and IC. In addition, elevated levels of DoxSL have been implicated in other diseases, such as type 2 diabetes, diabetic sensory neuropathy, von Gierke disease, and non-alcoholic hepatosteatosis [35]. Therefore, unraveling the cytotoxicity of DoxSL is key to understanding the pathogenesis of HSAN IA/C and the roles of DoxSL in the other diseases.

The earliest symptom and the clinical hallmark of HSAN IA/C is loss of pain and temperature sensations in the distal extremities [1]. Intense mechanical, chemical, and thermal stimuli on the extremities are sensed by free nerve endings named nociceptors in between skin cells. In nociception, the stimuli activate various ion channels in the plasma membranes of the nociceptors leading to action potentials that travel through the sensory neuron to the central nervous system [36–38]. We found that C_26_-DoxDHCer is more toxic to the cell than C_16_- or C_18_-DoxDHCer. It is possible that C_26_-DoxDHCer is the most toxic DoxSL species. In humans, very long-chain (C_22-26_) ceramides are synthesized by CerS3, which is highly expressed in skin cells [25]. Therefore, we hypothesize that (1) skin cells should be directly affected by elevated levels of DoxSL and that (2) unhealthy skin cells are contributors to the origin of this clinical hallmark and other clinical features of HSAN IA/C. The clinical hallmark raises several questions. (1) What causes loss of pain and temperature sensations? The importance of healthy skin cells for nociception is underscored by findings from studies on peripheral neuropathy as a side effect of paclitaxel in cancer chemotherapy. In mammalian cells, paclitaxel induces dose-dependent increase in the level of DoxSL [18]. In zebrafish, paclitaxel promotes epithelial damage and reduces mechanical stress resistance of the skin. These perturbations precede degeneration of sensory axons and loss of touch response in the distal caudal fin [39]. These findings indicate that unhealthy skin cells due to elevated levels of C_26_-DoxDHCer might lead to degeneration of sensory neurons innervating the skin and to loss of pain and temperature sensations in the extremities. (2) Why are sensory functions more affected than motor functions? Skeletal muscles mainly express CerS1 and 5 which normally produce C_18_- and C_16_-ceramides, respectively [25]. Since C_18_- or C_16_-DoxDHCer is less toxic than C_26_-DoxDHCer (at least in yeast), skeletal muscles and neuromuscular junctions would be less affected than skin cells and nociceptors in the skin. Therefore, sensory functions are more affected than motor functions of the extremities. (3) Why are the distal parts more affected than the proximal parts? The density of nociceptive innervation in the distal parts is less than that in the proximal parts of extremities [40]. Therefore, nociception in the distal parts is more sensitive to perturbations leading to reduction in the number of functional nociceptors than that in the proximal parts of the extremities.

Individuals with HSAN IA/C might suffer from disturbances related to other skin functions, such as hypohidrosis (diminished sweating) [1] and recurrent chronic skin ulcers that heal very slowly (up to a year). Such ulcers might develop from minor wounds that heal very slowly [1] or without an apparent cause [41]. These clinical features further indicate that skin cells are sensitive to elevated levels of DoxSL. Therefore, we also hypothesize that elevated levels of C_26_-DoxDHCer produced by CerS3 in skin cells perturb sweat glands in the skin and inhibit the renewal and accelerate the death of skin cells, thereby reducing sweat production and slowing down skin wound healing or causing skin lesions without an apparent cause, respectively.

There are eight missense mutations in *SPTLC1* and six missense mutations in *SPTLC2* that have been conclusively linked to HSAN IA and IC, respectively [42–44]. Different mutations increase the substrate promiscuity of SPT towards *L*-alanine and glycine to different degrees, leading to different accumulation rates of DoxSB [42]. We found that low accumulation rates of DoxSa allow slow accumulation of DoxDHCer for long periods of time until DoxDHCer reaches its toxic level, whereas high accumulation rates of DoxSa lead to brief periods of DoxDHCer production as DoxSa rapidly reaches its toxic level. We also found that the DoxDHCer species responsible for the toxicity of DoxDHCer changes over the course of its accumulation. The most toxic species (C_26_-DoxDHCer) is mainly responsible for the toxicity when the levels of DoxDHCer are low, whereas all species are responsible for the toxicity when the levels of DoxDHCer are high. Since individual DoxSB and DoxCer species might have different mechanisms of action, our findings suggest that (1) the mechanism of cytotoxicity might vary in affected individuals with different causing mutations and that (2) it might evolve over the course of the development of the disease.

Finally, this study highlights the importance of acyl chain length in the toxicity of DoxDHCer. Since individual DoxSB and DoxCer species might have different mechanisms of action, further studies should be focused on dissecting the toxicity of individual DoxSB and DoxCer species.

## Materials and Methods

### Yeast strains

**Tab 1.**
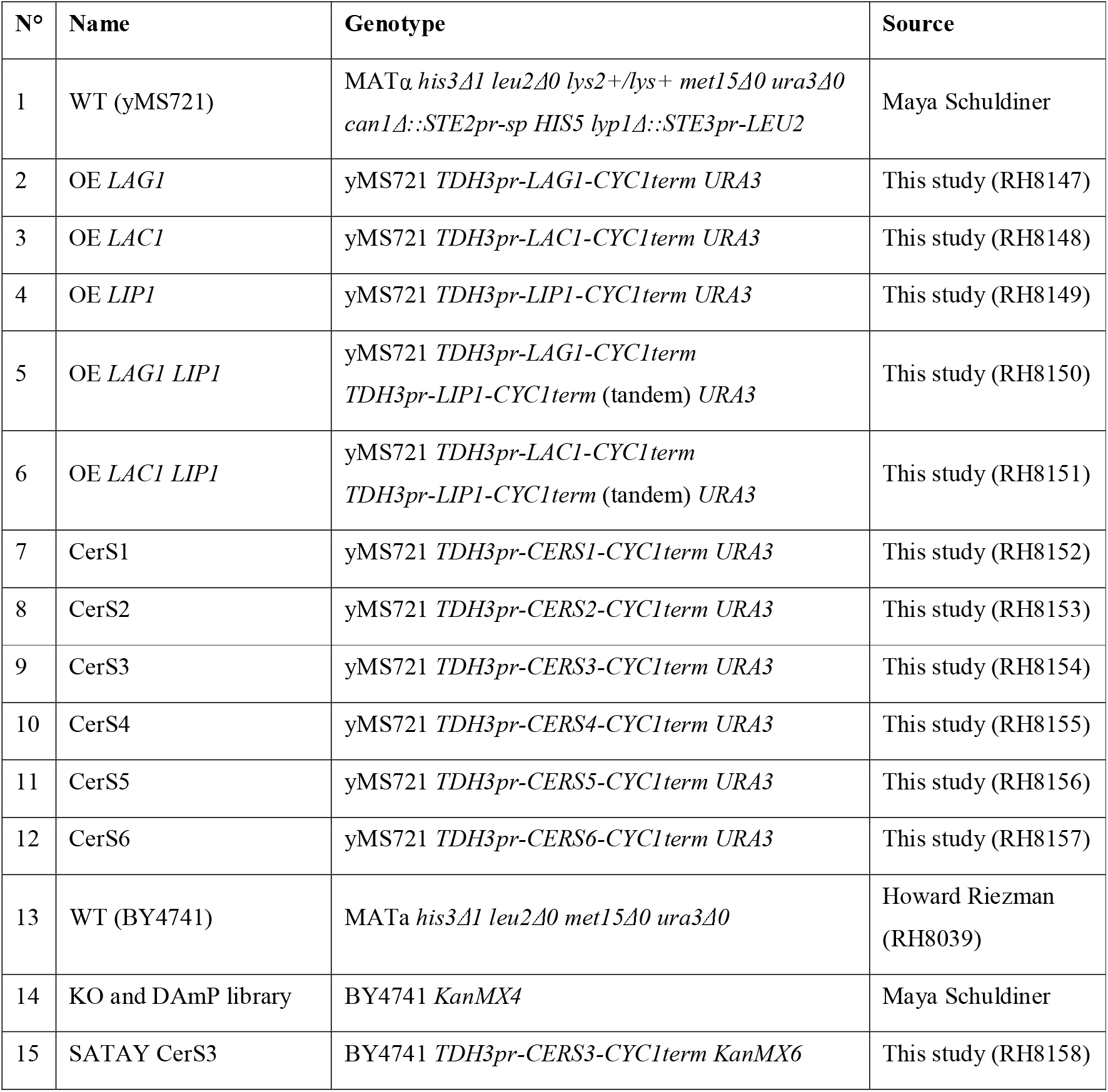
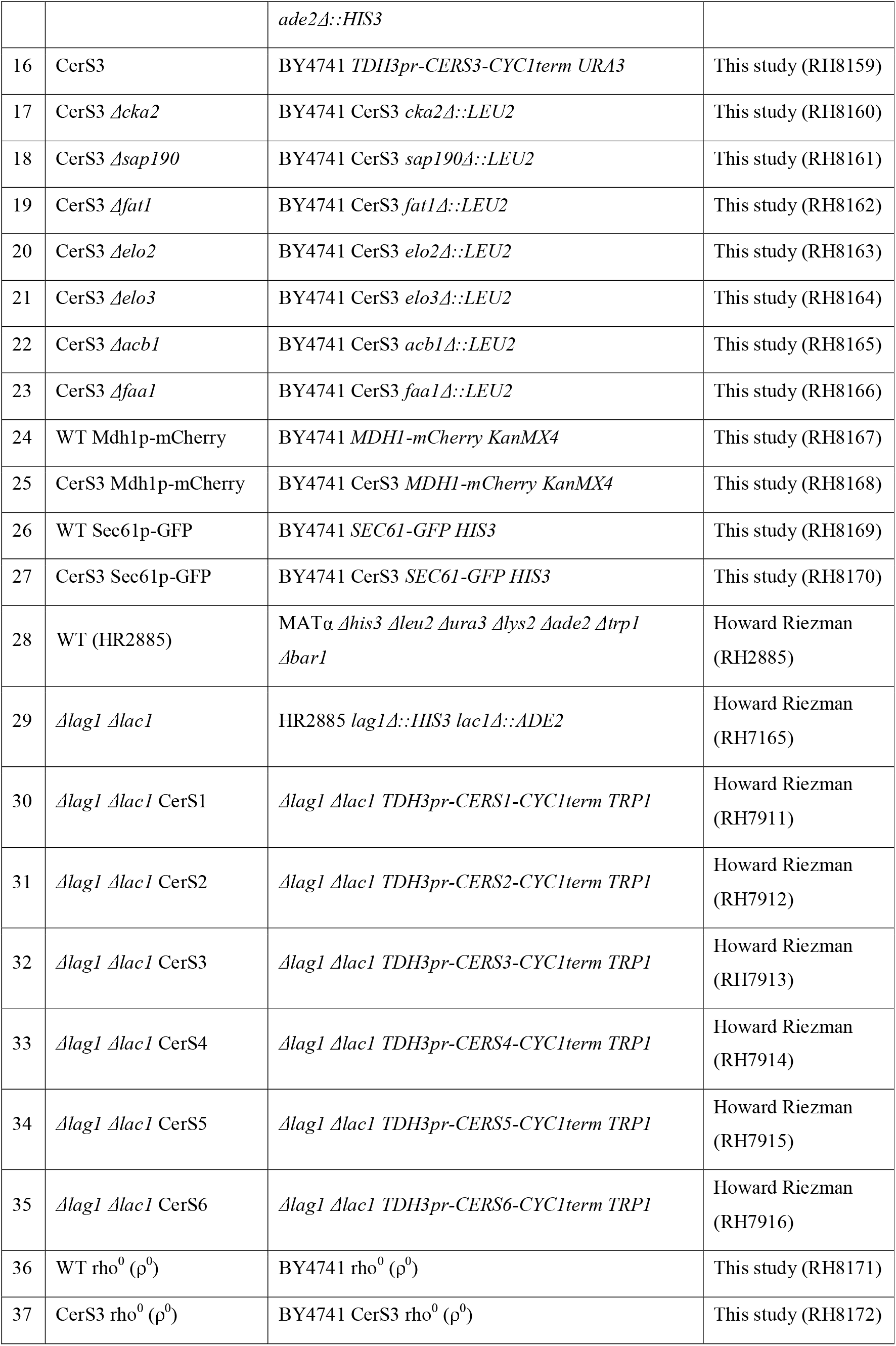
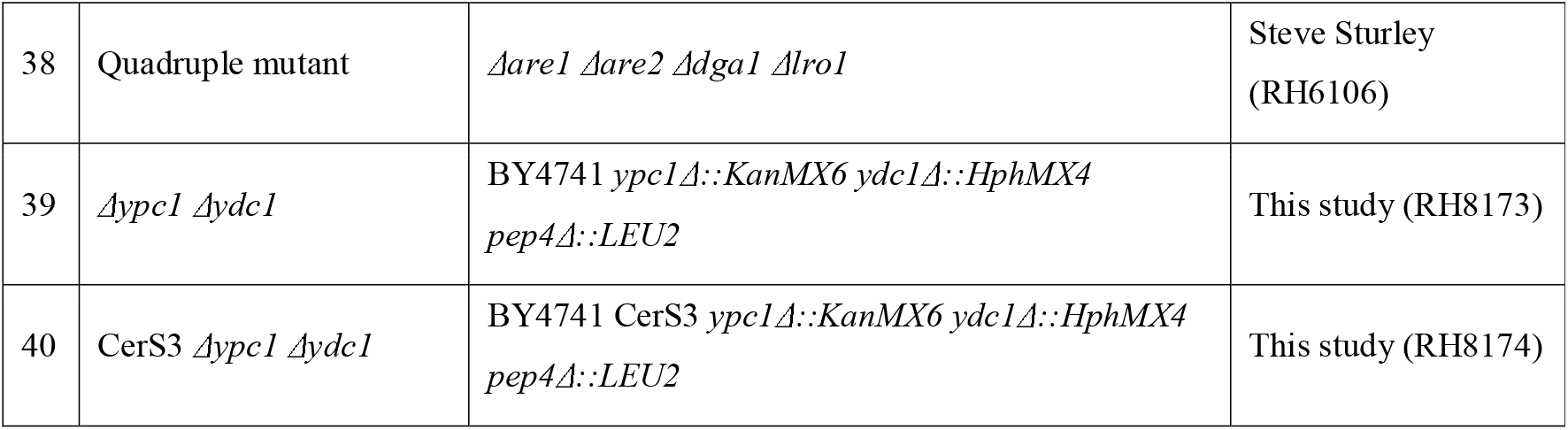
Yeast strains used in this study.

### Lipids

Sphinganine (Sa) (Avanti 860498), 1-deoxysphinganine (DoxSa) (Avanti 860493), 1-deoxymethylsphinganine (DoxmetSa) (Avanti 860473). The lipids were dissolved in ethanol and sonicated for 5 min using an ultrasonic bath to make stock solutions. The solutions were stored at −20°C. Before use, the solutions were brought to room temperature and sonicated for 5 min.

### Spot assay

YPD agar supplemented with 10 mM of MES, 0.05% (^v^/_v_) of Tergitol^TM^ NP-40, and a lipid was prepared two days in advance. On the spotting day, cells were suspended at a ten-fold serial dilution with OD_600_ of 1.5 at the highest in 200 μl of YPD liquid in a U-bottom 96-well plate. The cells were spotted on the agar medium using a 48-pin tool. The culture was incubated at 30°C for 2 days.

### Lipidomics analysis

#### DoxSa treatment

Exponentially growing cells (OD_600_ of 1 or 20 million cells/ml for non-toxic DoxSa treatment, or OD_600_ of 0.05 or 1 million cells/ml for the standardized DoxSa treatment) in YPD liquid were treated with 1-deoxysphinganine (DoxSa) at 30°C for the indicated durations. Then, the metabolism of the cells was immediately quenched with 5% of ice-cold trichloroacetic acid.

#### Lipid extraction

Lipid extraction was performed as described before [45] with minor modifications. Briefly, a mixture of lipid standards (7.5 nmol of 17:0/14:1 PC, 7.5 nmol of 17:0/14:1 PE, 6 nmol of 17:0/14:1 PI, 4 nmol of 17:0/14:1 PS, 1.2 nmol of C_17_-ceramide, 1.2 nmol of C_12_-1-deoxydihydroceramide, and 2 nmol of C_8_-glucosylceramide) was added to 25 OD_600_ values of cells. Then, the cells were subjected to two-round lipid extraction with 1.5 ml of extraction solvent (ethanol, water, diethyl ether, pyridine, 4.2 N of ammonium hydroxide 15:15:5:1:0.018) and 250 μl of glass beads by vigorous vortexing for 5 min followed by incubation at 60°C for 20 min. Cell debris was pelleted at 800 *×g* for 5 min, and the supernatant was collected. The combined supernatant was divided into two equal aliquots for glycerolipid and sphingolipid analyses. Both aliquots were dried with a stream of nitrogen. The sphingolipid aliquot was then treated with 0.5 ml of monomethylamine reagent (methanol, water, *n*-butanol, methylamine 4:3:1:5) at 53°C for 1 h and dried again. Next, both dried aliquots were subjected to three-round desalting by resuspending them in 300 μl of water-saturated *n*-butanol and 150 μl of water followed by centrifugation at 3,200 *×g* for 10 min to induce phase separation. The upper phases were collected. To start another round of desalting, the lower phase was mixed again with 300 μl of water-saturated *n*-butanol. The combined upper phases were dried and stored at −80°C.

#### Mass spectrometry (MS)

The dried lipid extracts were dissolved in 500 μl of chloroform:methanol (1:1). Each extract was diluted with chloroform:methanol:water (2:7:1) or chloroform:methanol (1:2) containing 5 mM of ammonium acetate for positive- or negative-mode MS, respectively. Then, the samples were infused into a TSQ Vantage mass spectrometer (ThermoFisher) using a Nanomate (Advion) with a gas pressure of 30 psi and a spray voltage of 1.2 kV for multiple reaction monitoring analyses. The mass spectrometer was operated with a spray voltage of 3.5 kV in positive mode and 3 kV in negative mode. The capillary temperature was set to 190°C. Lipid amounts were normalized by the amounts of inorganic phosphate or OD_600_ values of cells.

### Genome-wide genetic screen of knockout and hypomorphic mutants

#### Synthetic genetic array

The CerS3 strain (in yMS721 background) was used as a donor strain to introduce the gene into every strain in the knockout or hypomorphic (DAmP) allele library [46, 47] by the synthetic genetic array (SGA) method [26, 27]. Briefly, the CerS3 strain was pinned onto 1536-format arrays of mutant colonies on YPD agar using a pinning robot (Singer Instruments) and incubated at room temperature for 1 day to induce mating. Then, the cells were pinned onto SD MSG agar – Ura + 200 mg/ml of G418 and incubated at 30°C for 1 day to select for diploid cells. The selection was repeated once. Next, the cells were pinned onto SPO agar and incubated at room temperature for 5 days to induce sporulation. The plates were wrapped with a moist towel to prevent desiccation. Then, the cells were pinned onto SD MSG agar – His/Lys/Arg/Ura + 50 mg/l of canavanine + 50 mg/l of thialysine and incubated at 30°C for 2 days to select for cells with the same mating type as that of the mutants. Next, the cells were pinned onto SD MSG agar – His/Lys/Arg/Ura + 50 mg/l of canavanine + 50 mg/l of thialysine + 200 mg/ml of G418 and incubated at 30°C for 1 day to select for the mutants bearing CerS3. The selection was repeated once. The library was maintained on the same medium without G418.

#### DoxSa treatment and colony size measurement

The cells in the library were pinned onto SD MSG agar – His/Lys/Arg/Ura + 50 mg/l of canavanine + 50 mg/l of thialysine + 0.05% (^v^/_v_) of Tergitol^TM^ NP-40 supplemented with DoxSa and incubated at 30°C for 1 day. Then, the plates were scanned with a paper scanner (Hewlett Packard). The size of colonies was determined with the “Balony” software [48] by including the row-column correction.

### Genome-wide genetic screen of transposon-insertion mutants (SATAY)

#### Library generation

SAturated Transposon Analysis in Yeast (SATAY) was performed as described before [31] with minor modifications. Briefly, the SATAY CerS3 strain was transformed with pBK257 plasmid containing the transposon. Freshly transformed cells were inoculated into 1 l of SC liquid + 0.2% of glucose + 2% of raffinose – Ura at OD_600_ of 0.15 and grown at 30°C until saturation (OD_600_ of 3-4). Then, the culture was concentrated ten times by centrifugation at 600 *×g* for 5 min to obtain final OD_600_ of 37. To induce transposition, the cells were plated using glass beads onto 433 8.5-cm Petri dishes of SC agar + 2% of galactose – Ade and incubated at 30°C for 3 weeks. Contaminated plates were removed during the incubation time. Next, colonies were scrapped using a glass rod with minimum amounts of SC liquid + 2% of glucose – Ade, pooled, inoculated into 1 l of SC liquid + 2% of glucose – Ade at OD_600_ of 0.125, and incubated at 30°C until OD_600_ of 0.5. The library was used immediately.

#### DoxSa treatment

The cells in the library were pelleted at 800 *×g* for 5 min, inoculated into pre-warmed 500 ml of SC liquid + 2% of glucose – Ade supplemented with DoxSa at OD_600_ of 0.1, and incubated at 30°C until saturation. The treatment was repeated once. Next, the cells were harvested by centrifugation at 2,500 *×g* 4°C for 5 min and stored at −80°C.

#### DNA preparation

Genomic DNA of about 500 mg of cells was extracted by the phenol/chloroform extraction method. Next, 2 μg of genomic DNA was digested with 50 units of DpnII or NlaIII at 37°C for 24 h. The enzymes were then heat inactivated at 65°C for 20 min. The DNA fragments were circularized with 25 Weiss units of T4 Ligase at 22°C for 6 h. The circularized DNA molecules were precipitated with 0.3 M of sodium acetate pH 5.2, 1 ml of ethanol, and 5 μg linear acrylamide (Ambion AM9520) at −20°C overnight. Then, DNA was pelleted at 16,100 *×g* 4°C for 20 min, washed with 1 ml of 70% ethanol, and dried at 37°C for 10 min. Next, transposon fragments were amplified with Taq polymerase (New England Biolabs). The PCR products were then purified with the PCR clean-up/gel extraction kit (Macherey-Nagel) according to the manufacturer instruction, with the following modifications. DNA was bound to the column by centrifugation at 3,000 *×g* for 30 s. Then, 30 μl of elution buffer (10 mM Tris-HCl pH 8.5, 0.1% (^v^/_v_) Tween^®^ 20) was applied to the column, incubated for 3 min, and eluted by centrifugation at 11,000 *×g* 20°C for 1 min. The eluate was reapplied to the column and a second elution was performed under the same conditions.

#### Deep sequencing

Equal amounts of DNA from DpnII- and NlaIII-digested samples were pooled and sequenced using MiSeq v3 chemistry, according to the manufacturer instruction.

### Fluorescence microscopy

#### F-actin

Exponentially growing cells (OD_600_ of 0.05 or 1 million cells/ml) in YPD liquid were treated with DoxSa and incubated at 30°C for 1.5 h. Next, the cells were fixed by directly adding paraformaldehyde to the culture at a final concentration of 4%. Then, 1 OD_600_ value of cells were washed with 3 ml of washing buffer (0.1 M of Potassium phosphate pH 7.5, 1.2 M of sorbitol), stained with 50 μl of 1 μM of phalloidin-Atto488 (Sigma-Aldrich) at 4°C for 1 h in the dark, and observed by confocal microscopy. The severity of the F-actin phenotype was measured as the area of globular structures stained with phalloidin-Atto488.

#### Mitochondria

Exponentially growing cells expressing Mdh1p-mCherry in YPD liquid were treated with DoxSa and incubated at 30°C for 1.5 h. Next, the cells were observed by confocal microscopy immediately without fixation. The severity of the mitochondria phenotype was measured as the area of globular structures highlighted by Mdh1p-mCherry.

#### Hydrophobic bodies

Exponentially growing cells in YPD liquid were treated with DoxSa and incubated at 30°C for 1.5 h. Next, the cells were fixed by directly adding paraformaldehyde to the culture at a final concentration of 4%. Then, 1 OD_600_ value of cells were washed with 3 ml of the washing buffer, stained with 1 μl of 1 mg/ml of Nile Red at 4°C for 15 min in the dark, and observed by confocal microscopy. The size of a hydrophobic body was measured as the area of the structure stained with Nile Red.

#### The ER membrane

Exponentially growing cells expressing Sec61p-GFP in YPD liquid were treated with DoxSa and incubated at 30°C for 1.5 h. Next, the cells were observed by confocal microscopy immediately without fixation. The severity of the ER membrane phenotype was measured as the circularity of the ER membrane. The circularity of the ER membrane was calculated as 4*π*\*area/perimeter^2. A value of 1.0 indicates a perfect circle. As the value approaches 0.0, it indicates an increasingly elongated shape.

### Anaerobic culture

Cells were spotted onto an anaerobic (supplemented with 10 mg/l of ergosterol and 420 mg/l of Tween^®^ 80) YPD, YPEG, or YPL agar medium. The Petri dish was placed inside an airtight chamber (ThermoFisher) equipped with a pack of oxygen-consuming reagent (ThermoFisher) and oxygen color indicators (ThermoFisher). The chamber was closed immediately and incubated at 30°C for 3 days.

### Generating rho^0^ (ρ^0^) cells

Exponentially growing cells in SD liquid were treated with 25 μg/ml of ethidium bromide from OD_600_ of 0.1 until saturation (24 h). The step was repeated twice.

### Growth curve assay

Cells were suspended at OD_600_ of 0.1 in 200 μl of YPD liquid in a flat-bottom 96-well plate. The cover of the plate was replaced with a gas permeable seal (4titude 4ti-0516-96). The culture was incubated at 30°C with agitation for 24 h in a plate reader (Biotek^TM^ Synergy^TM^ H1) while OD_600_ of the culture was recorded every 10 min.

## Acknowledgements

AGH, MM, MS, and HR were supported by the EU ITN “Sphingonet”. AGH was supported by Institute of Genetics and Genomics of Geneva (iGE3). AHG, JTH, and HR was supported by the NCCR Chemical Biology and the Swiss National Science Foundation. MS is an incumbent of the Dr. Gilbert Omenn and Martha Darling Professorial Chair in Molecular Genetics. MS is also supported by a VW foundation LIFE grant. AHM and BK were supported by the European Research Council (Starting Grant 337906-OrgaNet).

## Author Contributions

AGH, JTH, and HR conceived the study; AGH performed most of the experiments and the analyses; JTH assisted AGH in lipidomics; AGH and MM performed the genetic screen of knockout and hypomorphic (DAmP) mutants under the supervision of MS; AGH, AHM, and BK performed the genetic screen of transposon-insertion mutants (SATAY); AGH and HR wrote the manuscript; All authors read the manuscript and commented on it.

