## Supplementary material for "Cytotoxicity of 1-deoxysphingolipid Unraveled by Genome-wide Genetic Screens and Lipidomics": Supp Legends

**S1 Fig. Mammalian ceramide synthase expressed in yeast is functional**

**(A)** Growth of the indicated strains on a rich agar medium. **(B-H)** Levels of different classes of ceramide and complex sphingolipid in the indicated strains without or with a non-toxic DoxSa treatment (2 μM of DoxSa, 20 million cells/ml, 1.5 h) determined by MS.

**S2 Fig. Library coverage and evaluation of genome-wide genetic screen of knockout and hypomorphic mutants**

**(A)** Library coverage of the screen. **(B)** Colony sizes of the CerS3 strain on the indicated plates, illustrating the degrees of spatial bias. **(C)** Distributions of colony sizes on individual plates following the indicated treatments, illustrating the degrees of plate bias. **(D)** Correlations of colony size ratios in the two replicates following the indicated DoxSa treatments, illustrating the repeatability of the screen. **(E)** Coefficient of variation (CV) of colony size ratio following the indicated DoxSa treatments, illustrating the repeatability of the screen. **(F)** Similarity of fitness rankings of the mutants following the indicated DoxSa treatments, illustrating the consistency of the results.

**S3 Fig. Library coverage and evaluation of genome-wide genetic screen of transposon-insertion mutants (SATAY)**

**(A)** Library coverage of the screen. **(B)** Correlations of inserted transposon numbers following the indicated treatments illustrating the degrees of the loss of library complexity.

**S4 Fig. Accumulated Levels of C_14-24_-DoxDHCer in the resistant mutants**

**(A-F)** Levels of the indicated species of DoxDHCer in the indicated strains without or with a non-toxic DoxSa treatment (2 μM of DoxSa, 20 million cells/ml, 1.5 h) determined by MS. The red dashed lines indicate the levels of DoxDHCer species in the CerS3 strain following the treatment.

**S5 Fig. DoxSL toxicity is independent of mitochondrial respiration**

**(A)** Overview of the setup of anaerobic cultures. **(B)** Effect of oxygen on the growth of WT and the CerS3 strain cultured on fermentable (glucose) or non-fermentable carbon source (ethanol glycerol or lactate). The strains cultured on non-fermentable carbon sources could not grow inside the chamber. They resumed their growth once re-oxygenated. The color indicators and the growth of the strains indicate that the chamber was oxygen-free. **(C)** Effect of oxygen on the sensitivity of WT and the CerS3 strain to DoxSa. **(D)** Effect of fermentable and non-fermentable carbon sources on the sensitivity of WT and the CerS3 strain to DoxSa. **(E)** Growth of the indicated strains on agar media with the indicated carbon sources. ρ^0^ strains were not able to grow on glycerol, indicating that the strains lacked mitochondrial DNA. **(F)** Fitness defects of the indicated strains following the indicated DoxSa treatments. Area under curves were calculated from 24-h growth curves obtained from a plate reader.

**S6 Fig. Formation of hydrophobic bodies is independent of neutral lipids**

**(A)** Lipid droplets in WT and a quadruple mutant lacking the abilities to synthesize triglycerides and sterol esters (*Δare1* *Δare2* *Δdga1* *Δlro1*) following the indicated durations of culture in a rich liquid medium. The quadruple mutant was not able to form lipid droplets, indicating that formation of lipid droplets requires neutral lipids. **(B)** Hydrophobic bodies in the quadruple mutant following the indicated DoxSa treatments. Lipid droplets and hydrophobic bodies were stained with Nile Red. Each image is a maximum intensity projection of a 2-μm slice. The scale bars are 2 μm.

**Table Captions**

**S1 Tab.** Fitness of knockout and hypomorphic mutants bearing CerS3

**S2 Tab.** Fitness of transposon-insertion mutants bearing CerS3
