## Supplementary figures and images for "Cytotoxicity of 1-deoxysphingolipid Unraveled by Genome-wide Genetic Screens and Lipidomics"

### Supp figure 1

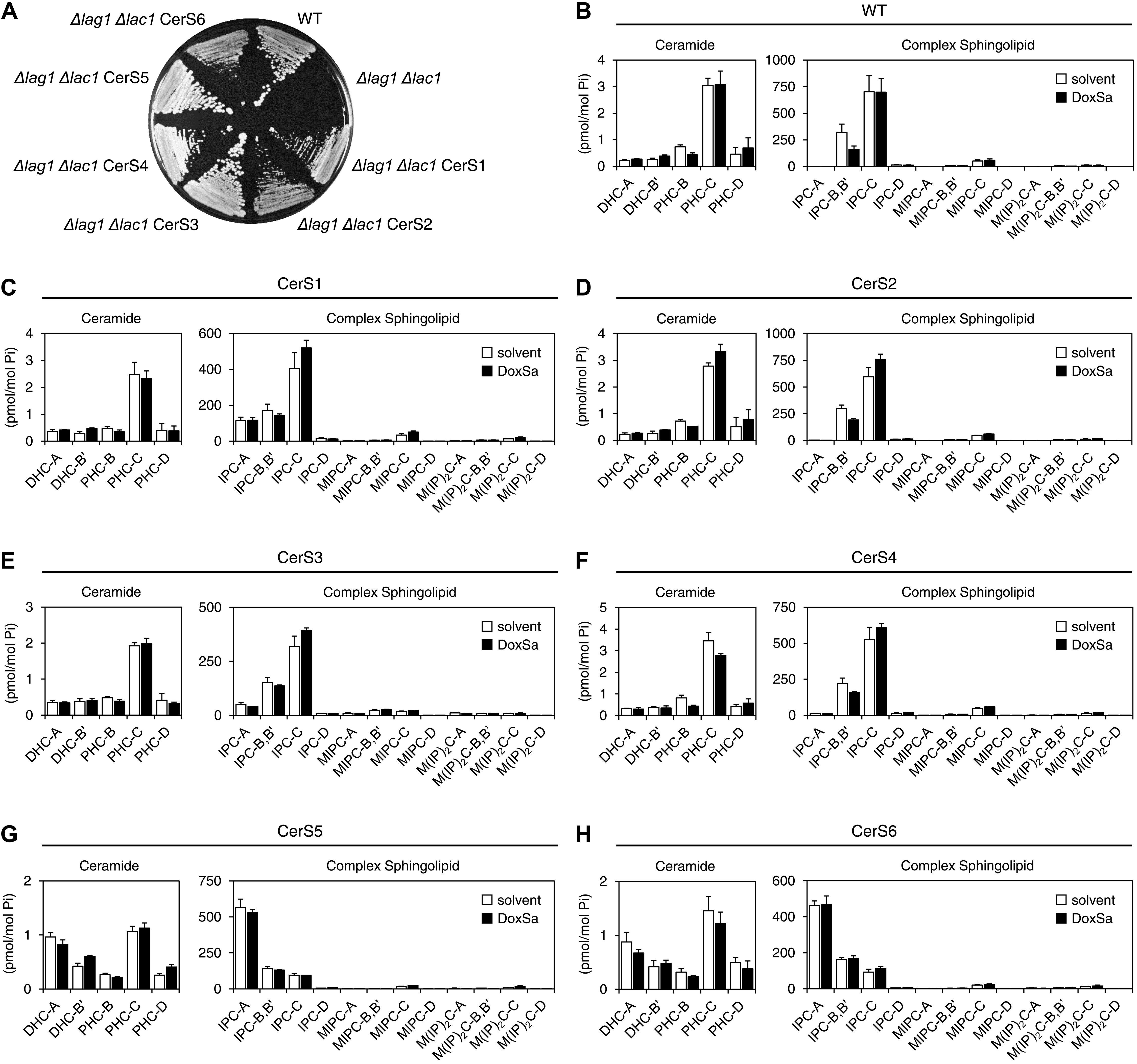

### supp figure 2

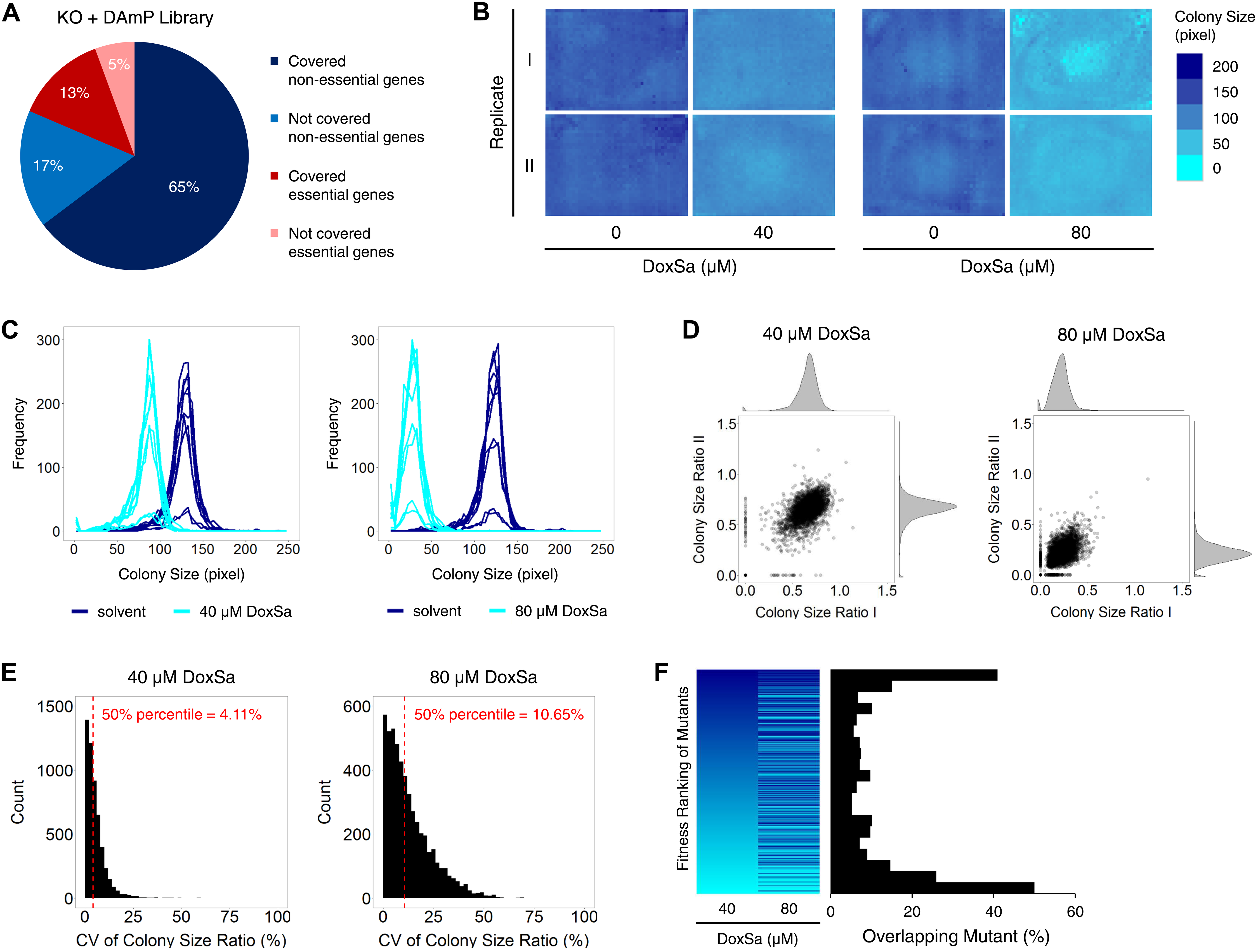

### supp figure 3

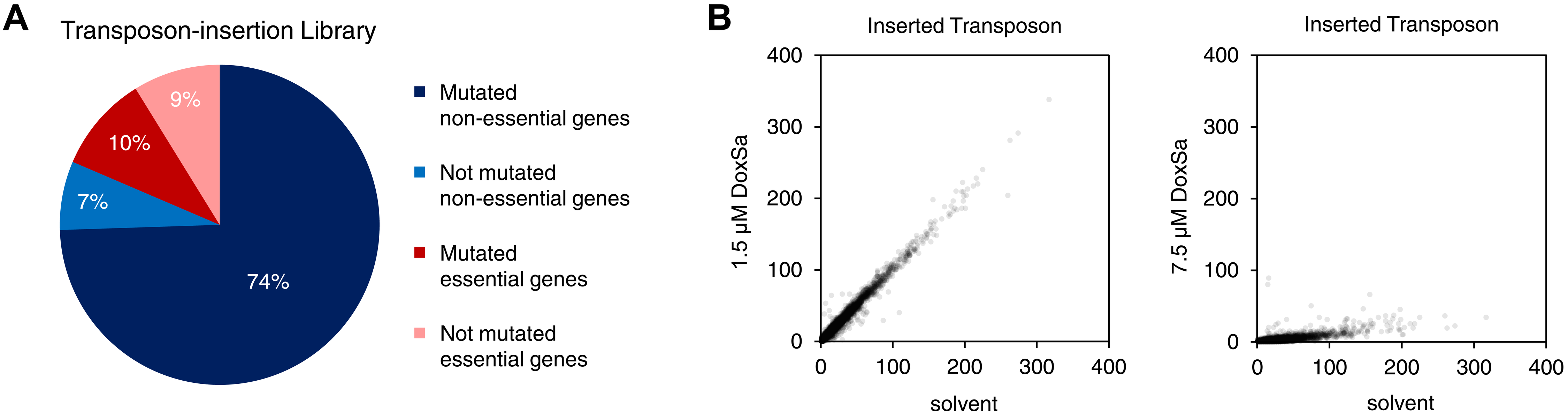

### supp figure 4

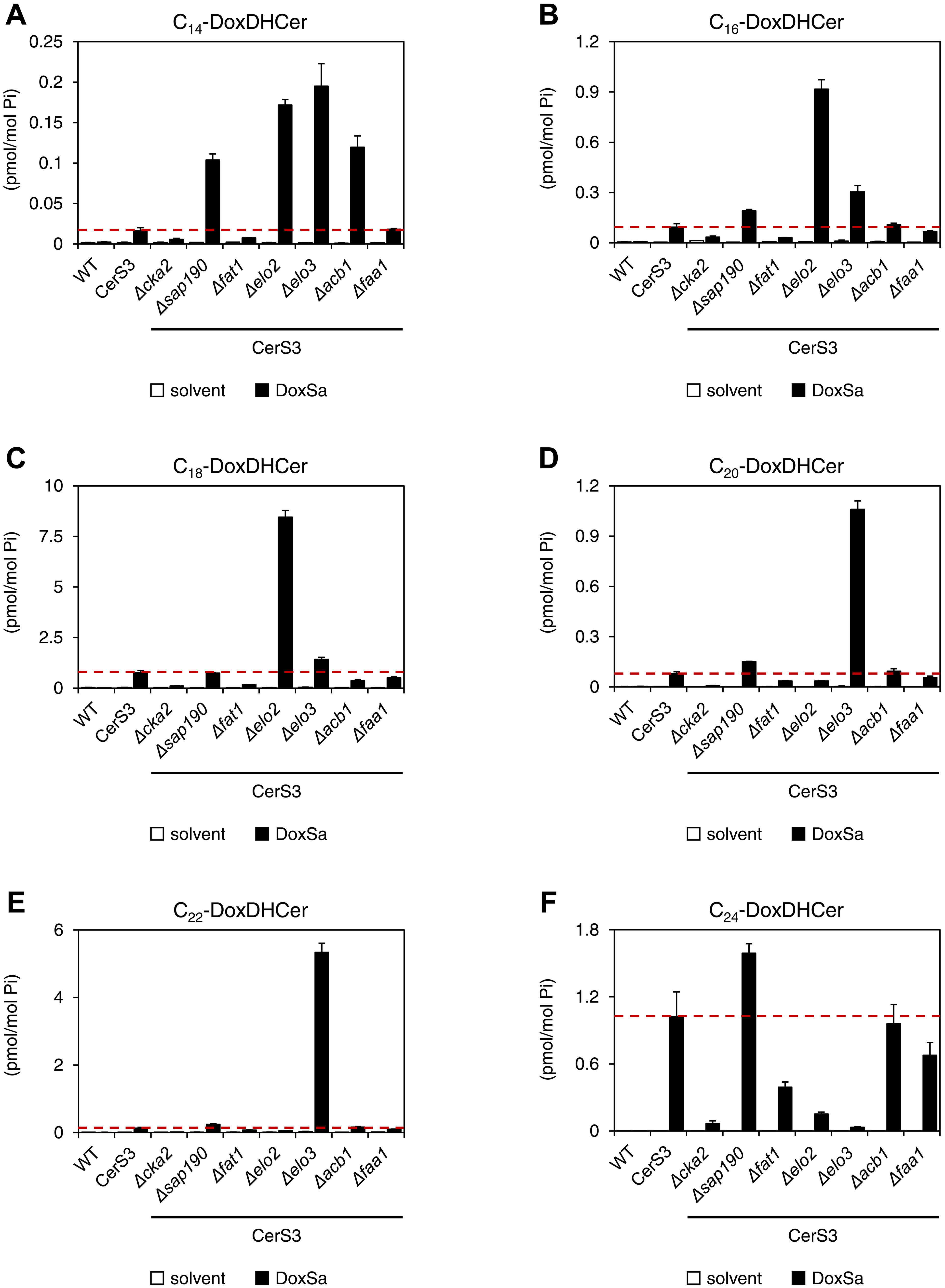

### supp figure 5

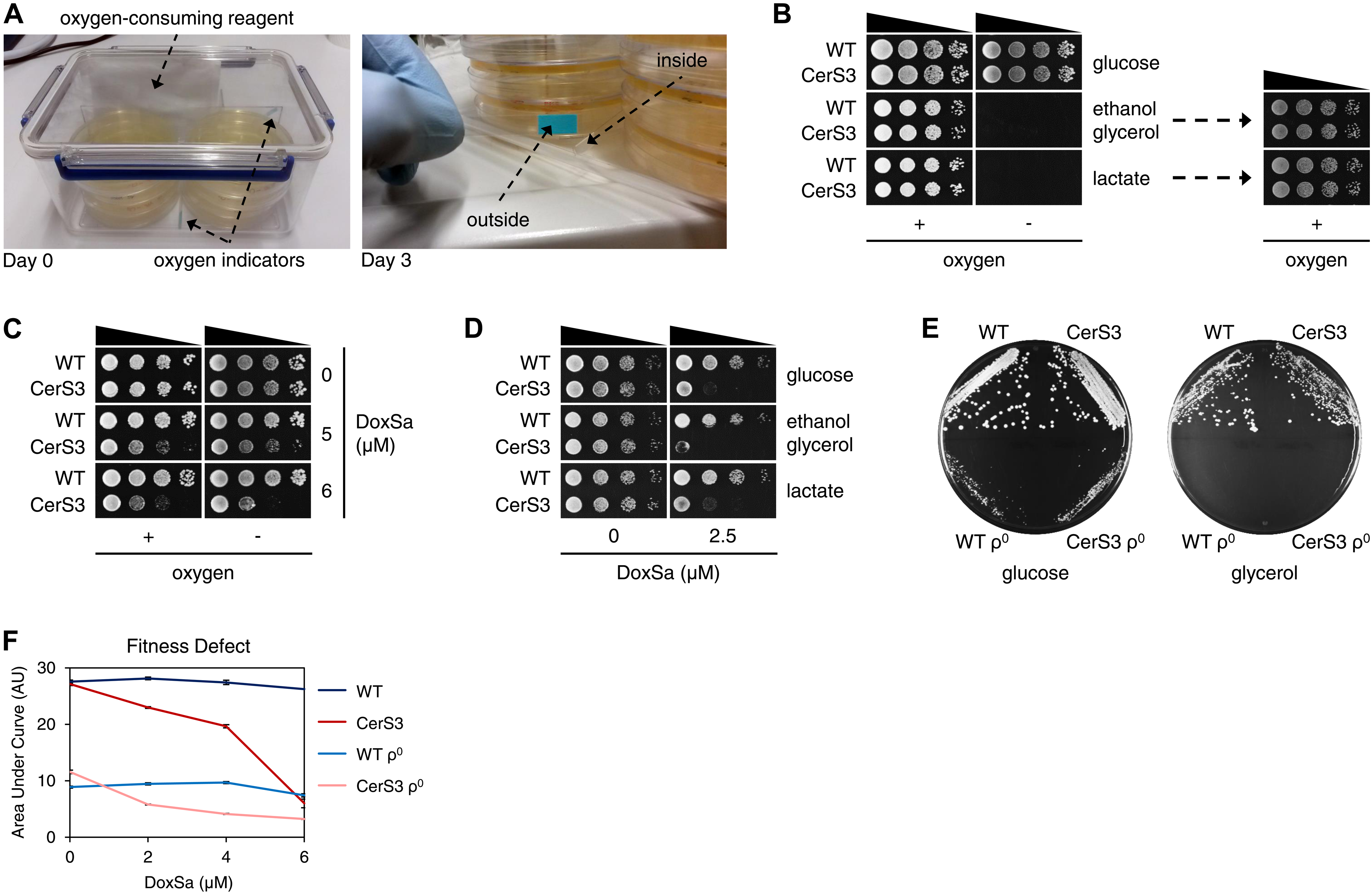

### supp figure 6

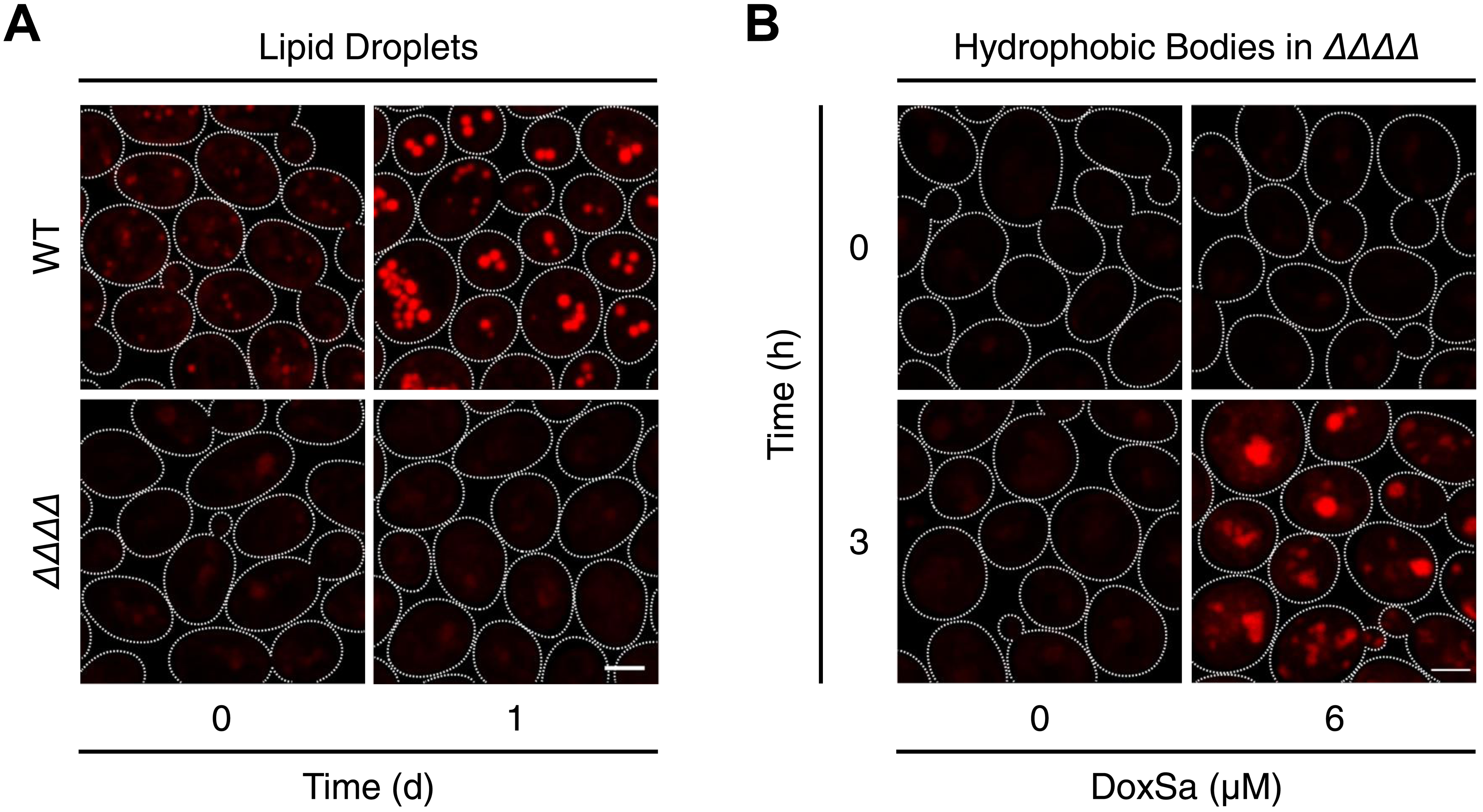
